# Drivers of intraspecies mitochondrial genome evolution revealed by deep population sequencing of a major fungal wheat pathogen

**DOI:** 10.64898/2026.11.27.769689

**Authors:** Ivan Skakov, Alice Feurtey, Pavel Kravchenko, Guido Puccetti, Ophélie Gning, Dmitry Knorre, Daniel Croll

## Abstract

Mitochondria play crucial roles in eukaryotic cells, serving as hubs for energy conversion. Despite functional conservation, mitochondrial genomes exhibit astonishing rates of expansion and rearrangement. Genome expansions correlate with the proliferation of selfish elements. However, how such variation arises in its earliest stages within species remains largely unknown. Here, we address this by establishing the largest intra-species survey of mitochondrial genomes focused on the major wheat pathogen *Zymoseptoria tritici*. We found that mitochondrial and nuclear population structures of 2193 genotypes were incongruent across the global distribution range. We systematically assembled mitochondrial genomes and determined major structural variants using combined short- and long-read sequencing. We found that 97.7% of all mitochondrial genomes cluster into seven haplotypes, distinguished mainly by two large insertions/deletions that underpin 11% variation in genome size. Recombination contributed to diversifying haplotypes. Selfish elements were absent, with the notable exception of a rare Eastern European haplotype encoding a GIY-YIG homing endonuclease. Mitochondrial genomes of closely related *Zymoseptoria* species revealed historical contingency in genome dynamics, with genome sizes ranging from 44-62 kb and variation in essential gene content. We traced the occurrence of the *Z. tritici* GIY-YIG homing endonuclease to a sparse and patchily distributed set of ascomycete mitochondria, consistent with horizontal transfer. Taken together, our study shows how changes in mitochondrial genome architecture among species can originate from extensive structural variation within species, and directs mechanistic studies on the impact of selfish element activity on genome evolution.

## Introduction

Mitochondria are vital eukaryotic organelles crucial for ATP production and various cellular functions ^1^ ^2^. Originating from endosymbiosis, mitochondria underwent a dynamic evolution of their genome with ongoing gene transfers to the nucleus ^3–5^. Mitochondrial genome size differs considerably among kingdoms, with *Metazoa* carrying mitochondria with compact genomes (typically 14-20 kb) ^6,7^, plant mitochondrial genomes ranging 180-600 kb ^8^, and fungi showing the widest range of mitochondrial genome sizes (18.8-332.2 kb) ^6,9^. Fungal mitochondrial genome variability is extensive even at the genus level. For instance, *Schizosaccharomyces* genome sizes vary 19-80 kb ^10^. Size variation is typically explained by the size of intergenic regions, variable GC clusters, tandem repeats ^11–15^, and most prominently by the variation in intron numbers and intron-encoded open reading frames (ORFs) ^12,13,16,17^. The number of introns in essential genes also shows extensive variation within single genera ^13,18–20^. For instance, in *Aspergillus* and *Penicillium*, multiple intron gain and loss events likely occurred ^19^. Self-splicing introns often include ORFs of homing endonucleases (HEs) ^21,22^, which are enzymes that recognize site-specific DNA targets. HEs can produce either double-strand breaks or single-stranded nicks, which lead to homologous recombination and enable selfish element replication ^23^. Fungal mitochondria typically encode GIY-YIG (GIY) and LAGLIDADG (LD) HEs, and rarely HNH endonucleases ^23^. Selfish element proliferation contributed to numerous, independent genome expansions across fungal clades ^24,25^. How such variability originates in individual species remains poorly investigated.

Intra-specific mitochondrial genome size variation was documented across multiple fungal phyla including Glomeromycota ^11^, Ascomycota (including Saccharomycetes ^26^ ^27^ and Sordariomycetes ^28^ ^16^ ^18^), and Basidiomycota ^29^. Consistent with observations for between-species differences, a major source of variation within species is introns ^16^. In *Cordyceps militaris,* group I introns were horizontally transferred from another fungal species, creating within-species presence-absence variation of introns ^16^. In *Saccharomyces cerevisiae,* introns show a patchy distribution in the *cob* and *cox1* genes, contributing to genome size variation of 7.8-20 kb ^30^. Similarly, intron insertions are present in the core mitogenome of the multi-host pathogen *Fusarium oxysporum* ^28^. Intergenic region variability contributes even more substantially to *S. cerevisiae* genome size plasticity (ranging 45.3-69.8 kb) ^30^. In *F. oxysporum,* long variable regions of 6–16 kb drive the main differences among mitogenomes ^28^. These long variable regions include tRNA loci and ORFs encoding homing endonuclease domains. How non-intronic size variation arises remains largely obscure; however, non-homologous recombination has been proposed as a mechanism. During mating, fungi often inherit mtDNA from both parents ^31–36^. The resulting heteroplasmic stage can lead to mitochondrial fusions^34,37,38^ and recombination. Signatures of recombination were reported in Basidiomycota ^39,40^ and Ascomycota ^37,41–45^, including *F. oxysporum*, where discordance in mitochondrial genotypes was observed ^28,46^.

Despite knowledge of molecular mechanisms creating mitochondrial genome diversity, the extent and distribution of such polymorphisms within species remain largely unknown. In large genomic surveys, such as those for the *F. oxysporum* species complex ^28^, a mismatch between the nuclear phylogeny and mitochondrial variants was reported. Meanwhile, predominantly clonal reproduction within the species and obscure evolutionary relations between the main nuclear clades make the interpretation of mitochondrial variant distribution challenging. A similar nuclear-mitochondrial mismatch was reported in *S. cerevisiae* ^30^; the complex population history of the species and comparatively undersampled wild yeast populations make hypothesis testing difficult. Domesticated strains have experienced multiple bottlenecks driven by artificial selection, which can distort the observed patterns of mitochondrial genetic diversity. One of the best-sampled fungi is the ascomycete *Zymoseptoria tritici,* causing Septoria tritici blotch in wheat. Originating from the Middle East alongside its wild grass-infecting sister species, *Z. tritici* likely coevolved with domesticated wheat. Population genomic analyses of worldwide populations ^47,48^ revealed that the pathogen’s genetic structure largely reflects the introduction and spread of wheat across continents, with the highest diversity found in Middle Eastern and North African populations. Mitochondrial diversity in *Z. tritici* may be shaped by selection driven by QoI fungicides, which target the mitochondrial cytochrome *bc1* gene, in which a key resistance mutation (G143A) has independently arisen multiple times since the 1990s ^49^. The mitochondrial genome is compact at 44 kb, circular, and lacks introns or homing endonucleases^50^. Population-level mitochondrial studies have mostly used RFLP or amplicon multilocus sequencing ^51,52^ and have focused on specific geographic areas such as Tunisia, Morocco, and North America ^53–55^.

Here, we established a large collection of over 2000 fully assembled mitochondrial genomes from the global distribution range of *Z. tritici*. We combined short- and long-read datasets to ascertain structural variants. We also assembled representative genomes from the sister species to connect structural variant dynamics and selfish element content. Finally, we retraced the evolutionary history of newly discovered selfish elements across ascomycetes.

## Results

### Incongruent population structures of nuclear and mitochondrial genomes

We analyzed a large short-read dataset of 2193 *Z. tritici* isolates from 50 countries and all major wheat-producing areas ^47,48^ to assess both mitochondrial and nuclear population structures (**Figure 1A**). These isolates were gathered from naturally infected fields and were previously used to characterize global patterns of historic gene flow ^47^. The dataset covers the pathogen’s center of origin (the Fertile Crescent in the Middle East), as well as regions where wheat and the pathogen were introduced most recently (*i.e.*, the Americas and Oceania). We followed a joint genotyping approach and generated raw variant calls mapped to a complete nuclear and mitochondrial reference genome. We detected 1237 SNPs segregating in the mitochondrial genomes, corresponding to 2.8% of the genome length. A total of 378 SNPs were biallelic and consistently genotyped across all isolates. We identified the G143A substitution in the *cytB* gene, which confers resistance to QoI fungicides. The resistance mutation is widespread in Central and Northern Europe and New Zealand (**Figure 1B**, **Supplementary Figure 1A, B**). The resistance mutation is at lower frequencies in Southern and Eastern Europe, and rare in North Africa and North America. The resistance allele was absent from the Middle East, South America, and Australia, consistent with the lack of QoI fungicide applications prior to the isolate sampling.

**Figure 1.**
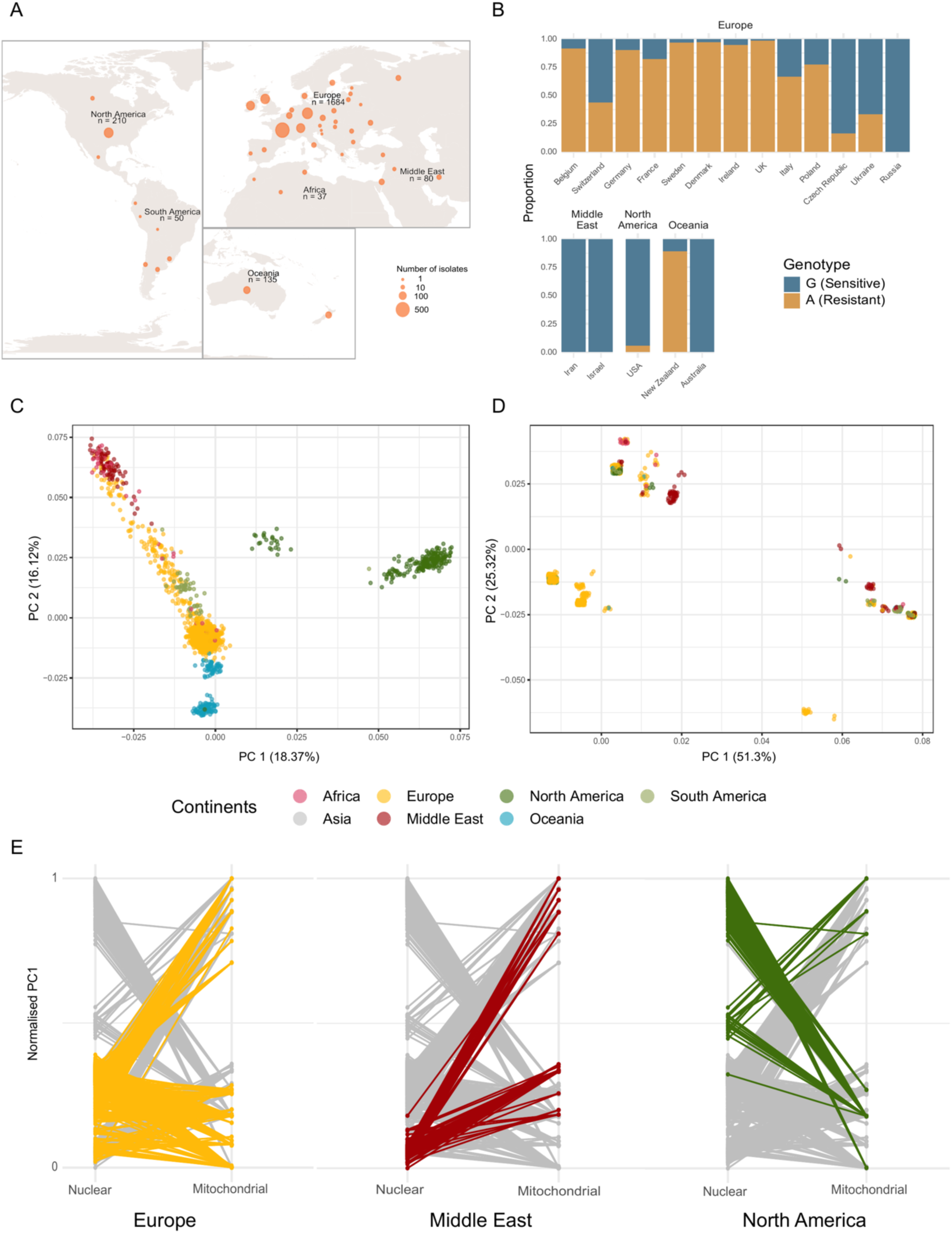
Comparison of the population structure defined based on the nuclear and mitochondrial genomes of *Z. tritici*. **A.** Map of the sampling scheme for the global collection of 2193 isolates of *Z. tritici*. **B.** Proportion of the isolates per country carrying the G143A mutation in the *cytB* gene, conferring resistance to QoI fungicides. Only countries with more than 20 sequenced isolates are shown. The G143A mutation is predominant in Europe and New Zealand, and absent or nearly absent in the Middle East, North America, and Australia. **C.** Principal component analysis (PCA), showing the first and second components (PCs) based on a subset of variants from the nuclear genome. Colors indicate the geographical origin of the isolates. **D.** PCA, showing the first and second components (PCs) based on variants from the mitochondrial genome. Coloring is identical to panel C. **E.** Discrepancy between nuclear and mitochondrial population structure: comparison of normalized PC1 coordinates for the nuclear (left axes) and mitochondrial (right axes) PCA analysis. The isolates from Europe, the Middle East, and North America are highlighted.

To assess discordance in nuclear and mitochondrial population structures, we first performed principal component analyses (PCA) separately for each genome. The PCA of the nuclear genome (based on 13,226 SNPs) revealed a nested structure with differentiation at the continental level and subdivisions within Europe and the Americas consistent with earlier reports ^47^ (**Figure 1C**; **Supplementary Figure 1C, D)**. In contrast, the PCA of mitochondrial genomes (based on 17 biallelic SNPs with a minor allele threshold of 2%) showed isolates falling into four main clusters without any apparent geographic association (**Figure 1D**; **Supplementary Figure 1E, F)**. To dissect the discordance, we constrasted the first principal components (PC1) of nuclear and mitochondrial genome PCAs for isolates from Europe, the Middle East, and North America (**Figure 1E**; **Supplementary Figure 1**). For European and Middle Eastern isolates, PC1 values were similar for nuclear genomes but clearly separated for mitochondrial SNPs. The opposite pattern was observed for North American isolates, with two distinct groups in the nuclear PC1, most of which clustered into a single group on the mitochondrial PC1.

To further investigate the mito-nuclear discordance patterns, we performed unsupervised sNMF-clustering analyses. For the nuclear genome, we identified 12 well-supported clusters (**Figure 2A**, **Supplementary Figure 2A–F**) consistent with previous analyses of global population structure ^47^. Given the higher number of European isolates included in our study (76%; 1680 out of 2193), we found evidence for an additional 12th cluster formed by Eastern European and Middle Eastern genotypes compared to previous analyses. A total of 86.7% (*n* = 1900) of nuclear genotypes were assigned to a single cluster (**Supplementary Figure 2A, B)**. The highest proportions of admixed genotypes were observed in Eastern European countries (86-100% admixed), Turkey (91%), and Italy (62%). These regions represent likely contact zones of distinct pathogen populations and hotspots of admixture. sNMF clustering applied to mitochondrial SNPs revealed three well-supported clusters (**Figure 2A**, **Supplementary Figure 3A–D**), which correspond to the groups observed in the mitochondrial PCA. Contrary to the nuclear clusters, the distribution of the mitochondrial clusters does not follow any evident continental-subcontinental differentiation (**Supplementary Figure 2B)**. To further investigate this, we constructed correspondence graphs between nuclear and mitochondrial clusters (**Figure 2B**, **C**). Isolates belonging to the North American nuclear cluster split between two mitochondrial clusters, while isolates of the nuclear North African cluster carry mitochondria assigned to any of the three mitochondrial clusters (**Figure 2B**). The largest mitochondrial cluster was represented by isolates covering all nuclear clusters, while the second-largest mitochondrial cluster was represented by isolates from 11 out of the 12 nuclear clusters (**Figure 2C**). Overall, we found strong evidence of mito-nuclear discordance across multiple regional clusters.

**Figure 2.**
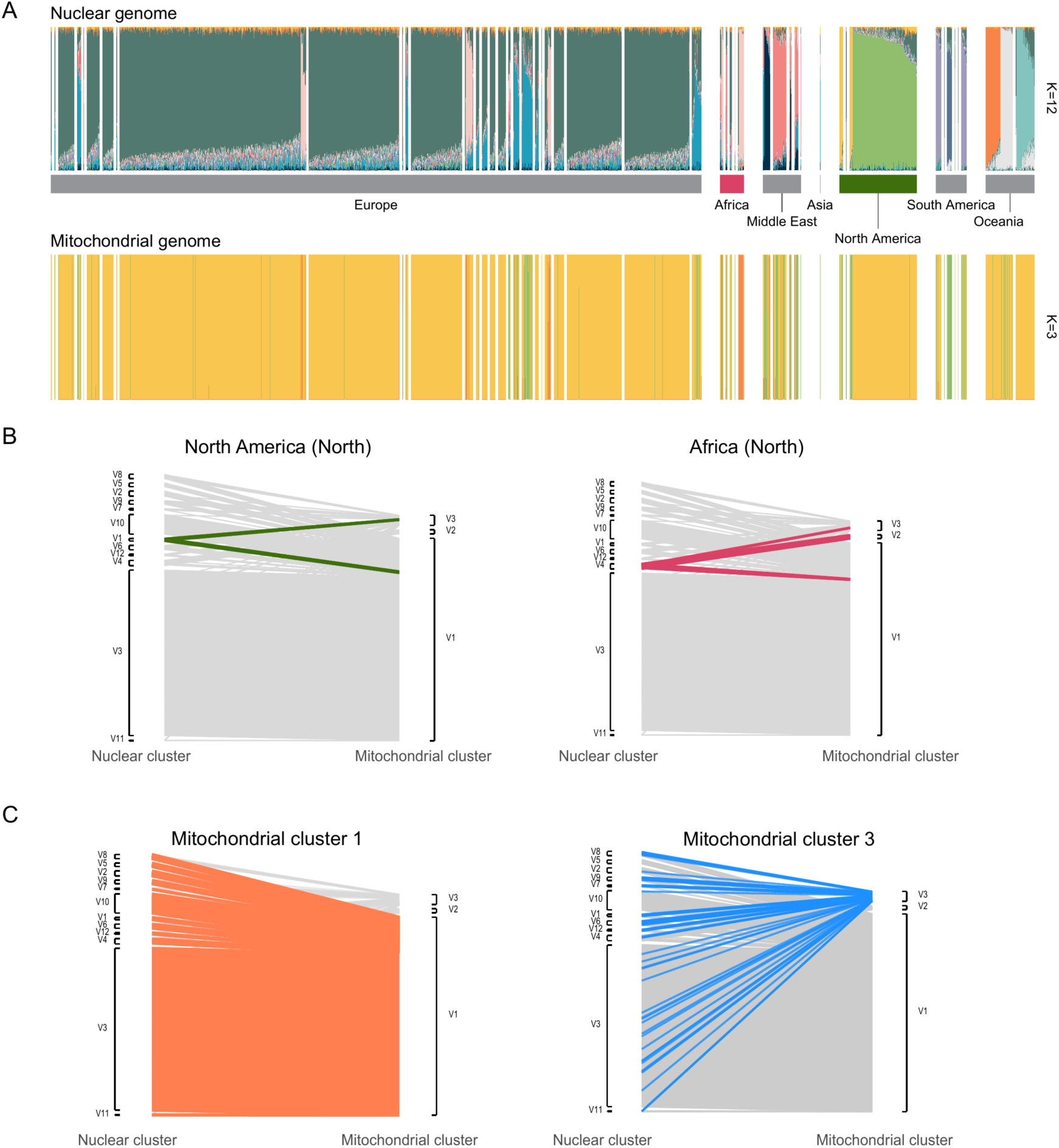
Comparison of the distribution of the isolates across the sNMF clusters defined for the nuclear and mitochondrial genomes. **A.** sNMF admixture coefficient plot for the nuclear (k=12) and mitochondrial genomes (k=3). **B.** Comparison of the assignments for the nuclear and mitochondrial sNMF clusters for the isolates from North America and Africa, highlighted by the green and purple colors, respectively. **C.** Distribution of the isolates from the clusters V1 (red) and V3 (blue) across nuclear clusters.

Discordance could potentially arise through an intermediary of mitochondrial heteroplasmy. To assess evidence for this, we compared the minor allele frequency (MAF) of the same sets of filtered biallelic SNPs in the nuclear and mitochondrial genomes. For each isolate, we calculated the mean proportion of reads supporting the minor allele across all polymorphic sites separately for the nuclear and mitochondrial genomes. As a reference, we used genotyping data of the haploid nuclear genome where only single alleles should be detected. Any low-frequency detection of a secondary base may have arisen from ambiguous mapping or unfiltered sequencing errors. We used this as a baseline to assess detection of a secondary allele in the mitochondrial genome. We found that the nuclear MAF was generally lower than mitochondrial MAF and highest MAF values exceeded the nuclear estimate by more than an order of magnitude **(Supplementary Figure 4A)**. Upon inspection of these isolates, we found that high-frequency variants were clustered into discrete regions of the mitochondrial genome **(Supplementary Figure 4B)**. Allele frequency distributions were largely bimodal unlike expectations from random sequencing or mapping errors. These patterns are consistent with the presence of at least one additional mitochondrial haplotype in some isolates **(Supplementary Figure 4C)**. However, a systematic investigation of evidence for heteroplasmy will require high-depth long-read sequencing.

### Intra-specific structural variation among mitochondrial genomes

To evaluate structural variation in mitochondrial genomes, we analyzed both short- and long-read datasets. A total of 23 isolates were sequenced with PacBio, representing the global diversity of the species and producing complete mitochondrial genomes 43.9-49 kb in length. A pangenome graph revealed variation defined by 348 variable positions (128 SNPs, 216 short indels, and 4 structural variants including indels > 30 bp; **Supplementary Figure 3**). All structural variants co-localized to two loci (positions 20,918 and 29,580 bp; **Figure 3A**, **B; Supplementary Figure 5A, B**), hereafter called rearrangement hotspots. The predominant haplotype was characterized by an insertion of 1774 bp in the first rearrangement hotspot, between the genes *nad2* and *atp9* (*n* = 12 isolates with haplotype 2; **Figure 3A-C**, **Supplementary Figure 5B, Supplementary Figure 6A, Supplementary Table 2**). Interestingly, the insertion is flanked by repeated sequences homologous to the beginning of the *nad2* gene (**Figure 3C**). Moreover, the 156-aa-long ORF predicted within the insertion retained homology to the upstream *nad2* gene. These features are consistent with a possible duplication of part of the *nad2* sequence. In the same rearrangement hotspot, we detected two isolates with a nested insertion of 3026 bp (**Figure 3A, B**, **C**, H**aplotype 3, Supplementary Figure 5B**). We detected three repeats within this insertion, with the second repeat being shorter than the other two (**Figure 3C**). The part of this insertion between repeats A and B is highly similar to the insertion in haplotype 2 described above, encoding the same fragmented 156-aa *nad2* copy, whereas the two other ORFs were not preserved. Between the repeats B and C, we identified a unique 2634 bp segment containing two ORFs: a short 48 aa ORF, homologous to the *nad2* gene, and a 795 aa ORF encoding the COX1 domain. The 85 aa close to the beginning of the last ORF are homologous to the cox1 gene located 16 kb upstream. The similarity in the structure of the flanking repeats suggests that the first part of the haplotype 3 insertion may have originated from duplication of the *nad2* region. However, the origin of the second part of the insertion remains unclear.

**Figure 3.**
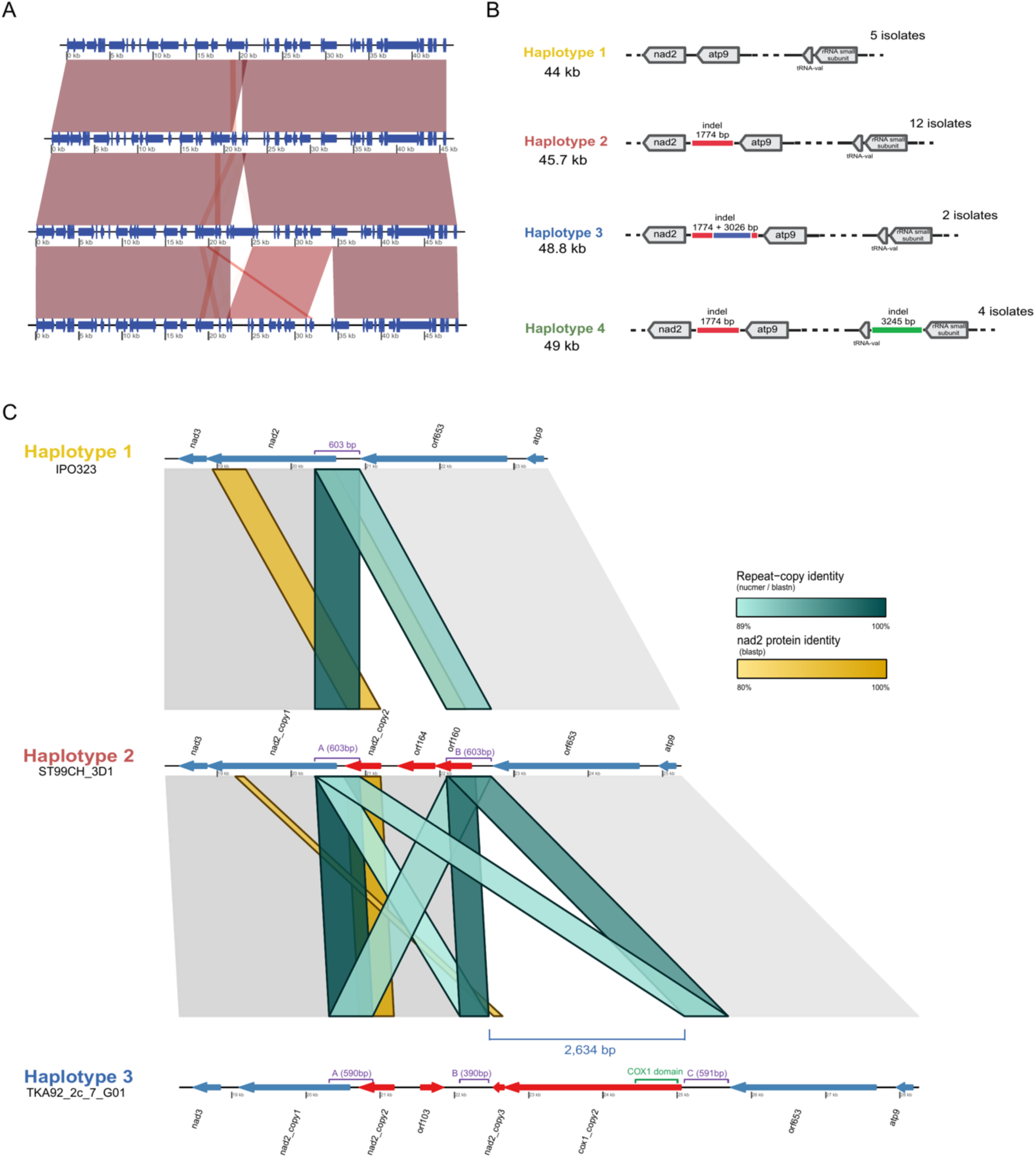
Reference long-read mitogenome assemblies. **A.** Synteny plot of the main 4 mitochondrial haplotypes, defined for 23 PacBio-sequenced isolates. Two main insertion/deletion points are clearly seen: in the 21 and 30 kb regions of the reference mitogenome. Short syntenic regions can be seen close to the 21 kb indel. **B.** Schematic representation of the insertions/deletion regions of the 4 main haplotypes, with the gene context of the variable regions. For each haplotype, the average length of the mitochondrial haplotype and the number of isolates sharing this haplotype among 23 isolates sequenced by PacBio are presented. **C.** The first rearrangement hotspot across three mitochondrial haplotypes: haplotype 1 carries a single ancestral copy of the 590–603 bp direct repeat and no insertion; haplotype 2 carries the short indel with two repeat copies (A, B) flanking a 1,171 bp insertion; haplotype 3 carries the long allele with three repeat copies (A, B, C) flanking a larger, 4,196 bp insertion. Protein-coding genes surrounding the insertions are depicted in blue, tRNAs are grey, and genes carried by the insertions are shown in red. The direct-repeat copies themselves are marked with purple brackets giving each copy’s length. The green bracket on cox1_copy2 marks the boundaries of its confirmed COX1 Pfam domain; the gold bracket beneath Haplotype 3 gives the size of the segment separating repeats B and C. Grey ribbons between rows show core flanking synteny from whole-genome nucmer alignment. Cyan ribbons show pairwise nucleotide %identity between every direct-repeat copy across all three haplotypes. Gold ribbons show blastp protein %identity of the nad2-derived ORFs.

An insertion of 3245 bp was identified in the second rearrangement hotspot, between the valine tRNA and small subunit rRNA genes (**Figure 3A**, **B, Haplotype 4, Supplementary Figure 5B, C**). The insertion is shared by four isolates and co-occurs with the 1767 bp insertion identifying haplotype 2. The insertion encodes two ORFs. The first 171 aa ORF encodes a protein with strong homology to *nad2* (**Supplementary Figure 5C**), whereas no conserved domains or homologous regions were identified for the second ORF, nor is the insertion flanked by repeats.

The long-read mitogenome assemblies provide high-confidence anchors to comprehensively map structural variation within the species. We generated mitochondrial draft genome assemblies for all 2,193 isolates with short-read sequencing data. We retained only circularized, single-contig assemblies ranging 42-55 kb (**Supplementary Figure 6B**), resulting in 1,973 assemblies, and excluded an additional 19 assemblies with rare and unusually large structural variants (6-24 kb) lacking support from long-read genome assemblies. We constructed a joint pangenome graph integrating both PacBio- and Illumina-based genome assemblies (*n* = 1977, **Figure 4A**). The graph comprised 1941 variants, including 1178 SNPs, 745 short indels, and 18 structural variants (>30 bp). The pangenome was represented overwhelmingly by haplotypes defined by singleton variants (933 out of 1099 haplotypes; 85%, **Supplementary Figure 6D**). Three haplotypes were shared by ≥50 isolates each. We identified two major hotspots of structural variation that define seven main haplotypes and account for 97.7% of all isolates. Four out of the five predominant haplotypes (**Supplementary Figure 6C**) match haplotypes found in long-read genomes and show indel variants of 1.7 and 3 kb at the first rearrangement hotspot (**Figure 4B**) and 3.3 kb at the second hotspot (**Figure 4C**). The first rearrangement hotspot is flanked by repetitive sequences. One of the five predominant haplotypes (haplotype 10, *n* = 41 isolates) was found exclusively in short read assembled genomes and was characterized by a partial translocation of the insertion from the first rearrangement hotspot to the second. We validated the presence of the 1.7 kb insertion by PCR, confirming that the insertion represents a genuine structural variant rather than an assembly artifact (**Figure 4D** **1, 2, Supplementary Figure 6E**).

**Figure 4.**
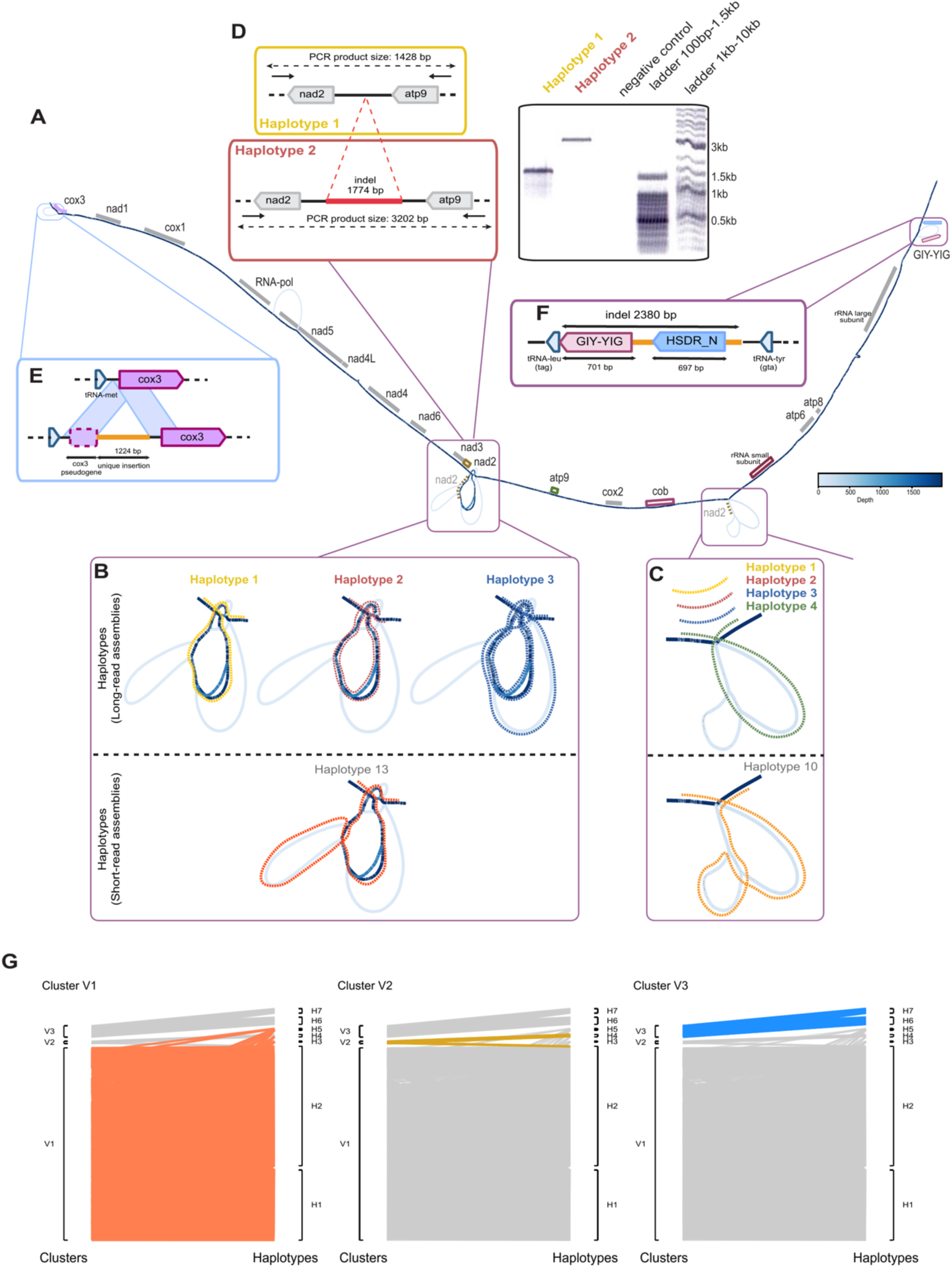
Diversity of mitogenomes within Z. tritici across 1,977 Illumina- and PacBio-sequenced isolates. **A.** Pangenome graph for 23 PacBio- and 1,954 Illumina-sequenced isolates, based on the multiple-sequence genome alignment. Shades of blue represent node coverage. The positions of essential mitochondrial genes are aligned with the pangenome. Two main structural variation hotspots are shown in panels **B** and **C**. Partial synteny to the *nad2* gene within the structural variants is represented by dashed yellow lines. The pangenome structure of the main PacBio haplotypes (1–4) is shown in the upper parts of the panels, while haplotypes identified only among Illumina-sequenced isolates are shown in the lower parts. Haplotypes are represented by dashed paths through the pangenome. **B.** First structural variation hotspot, showing indel variants of 1.7 and 3.3 kb. **C.** Second structural variation hotspot, showing a 3.2 kb indel. **D.** PCR validation of the 1.7 kb insertion defining PacBio haplotype 2. The first section shows the expected amplicon sizes for isolates carrying (ST95CH_3B2a, haplotype 2) and lacking (IPO323, haplotype 1) the insertion. The second section shows the corresponding PCR products. **E.** Haplotype 3, identified in 16 isolates, carries an insertion within the *cox3* coding sequence. The insertion contains a unique 520-aa ORF with cox3 and FACON_2 domains. **F.** The 2,380 bp insertion contains a GIY-YIG homing endonuclease identified in six isolates from Eastern Europe (Russia and Ukraine). The insertion is located between the *tRNA-leu* and *tRNA-tyr* genes and consists of a 701 bp GIY-YIG homing endonuclease-coding region, as well as the 697 bp gene with HSDR_N domain. The insertion is highly conserved, with one SNP among the six isolates carrying the insertion in the GIY-YIG gene and no SNPs in the HSDR_N gene and non-coding regions. **G.** Correspondence between mitogenome haplotypes defined by structural variation and mitogenome clusters defined by SNPs.

Structural variation in the pangenome graph was dominated by repetitive sequences originating through duplications and inversions (**Figure 4B**, **C**). Haplotype 3 (*n* = 16 isolates) carries an insertion in the coding sequence of the *cox3* gene without disrupting the gene, and encoding a 520 aa ORF with COX3 and BACON_2 domains (**Figure 4E**, **Supplementary Figure 6F, G**). Flanking repeats suggest that duplications are at the origin of this structural rearrangement. Finally, 6 isolates (∼0.4%) originating from Eastern Europe (Russia and Ukraine) carry a 2380- bp insertion located between the tRNA-leu and tRNA-tyr genes (**Figure 4F**). The insertion encodes a 233-aa ORF with a GIY-YIG homing endonuclease domain, as well as a 232 aa ORF with a type I restriction enzyme R protein N-terminus domain (HSDR_N, PFAM 13588). The vast diversity in structural variation captured by the pangenome graph expands the SNP-based clustering (**Figure 4G**). Distinct haplotypes captured by the pangenome graph were assigned to the same SNP-based cluster (e.g., haplotypes 6 and 7 both fall into cluster 3). The complexity of the intra-specific structural variation suggests ongoing generation of new variants and long-term retention.

### Patterns of recombination among mitochondrial genomes

Structural variation can arise from non-homologous recombination between different mitochondrial genome variants. To investigate evidence for recombination, we analyzed linkage disequilibrium among mitochondrial SNPs. We found a rapid decay of *r*² and |D’| (the normalized deviation from linkage equilibrium) in the recovered biallelic SNP set (**Figure 5A**, **B)**. However, SNP density was heterogeneous across the mitochondrial genome with notable enrichment near specific genic regions (**Figure 5C-D**). The two SNP-dense regions lie *ca.* 20 kb apart, i.e., roughly half of the circular genome, and are major contributors to the observed *r*² decay (**Figure 5C-D**). Evidence for linkage disequilibrium decay was robust to higher thresholds for minor allele frequency (**Figure 5A**). The decline in *r*² within 1 kb is consistent with at least sporadic recombination. We indeed detected recombination signals using multiple complementary tests (**Supplementary Figure 7A-F** and analysis of reticulation patterns (**Supplementary Figure 7G**). We reconstructed putative parental haplotypes and progenitors, revealing two large, repeated segments originating from one of the putative parents (**Figure 5** **D**). Additional segments located in a large insertion relative to the reference mitochondrial sequence suggest complex inheritance patterns (**Figure 5D**). A second putative parental genotype carried a segment encoding the *cytB* gene (**Figure 5D-E**). The substitution underpinning QoI resistance (G143A) is located within one of the recombining blocks (parent 4; **Figure 5D**), consistent with its propagation through mtDNA recombination. Previous evidence suggested that the resistance mutation arose in independent genetic backgrounds since the 1990s ^49^, however, evidence for a recombinant block comprising G143A means that a single origin is also plausible. Hence, mitochondrial recombination would have facilitated the spread of a major resistance factor.

**Figure 5.**
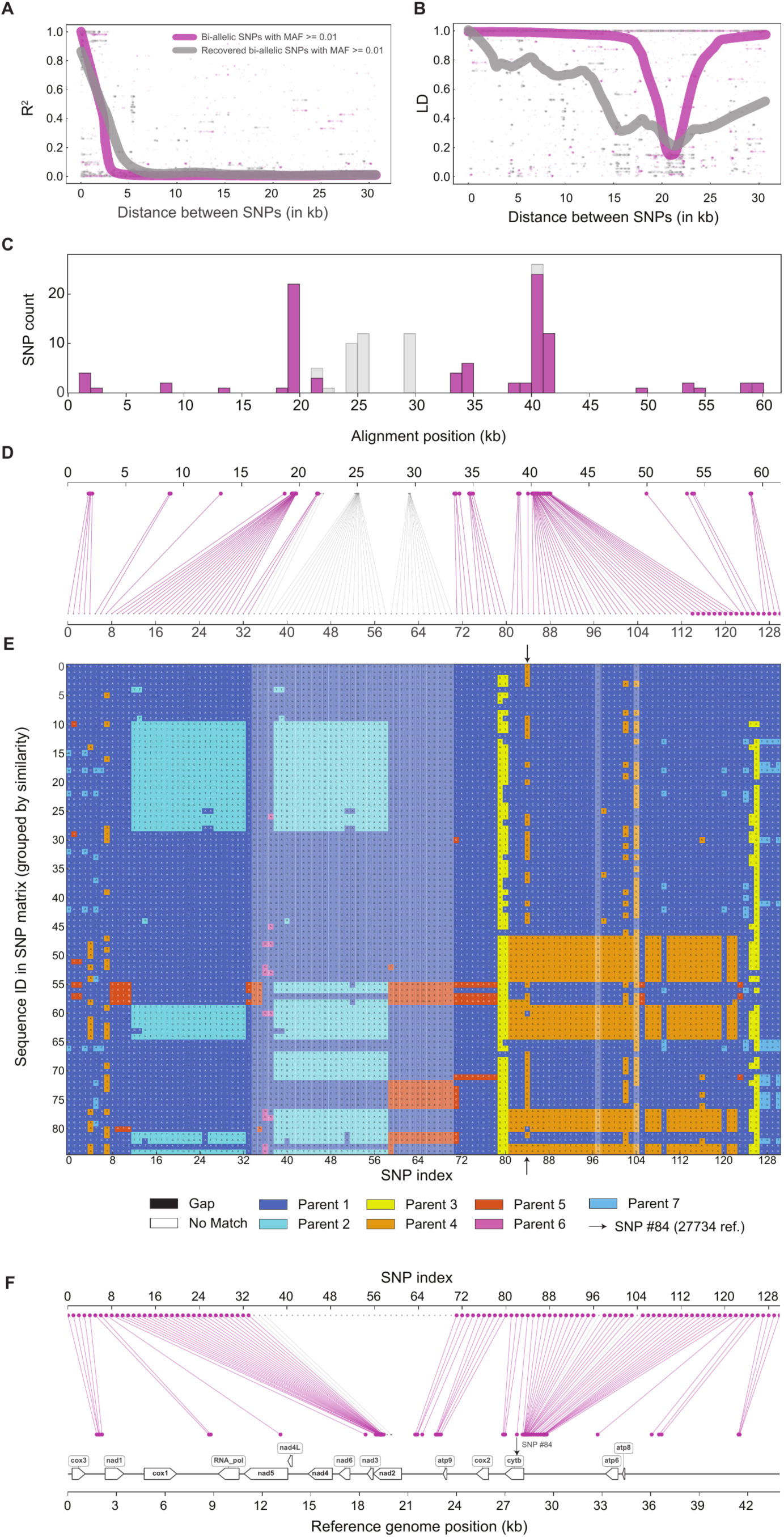
Linkage disequilibrium decay and population-wide SNP analysis suggest the presence of mtDNA recombination signal in the *Z. tritici* population. **A and B.** Linkage disequilibrium (R², |D’|) as a function of the distance between SNPs. R² and |D’| for the pairs of 111 phased biallelic SNPs that passed minor allele frequency filtration (MAF ≥ 0.01) are shown in gray. R² and |D’| for the extended set of recovered 131 SNPs (111 + 20 added from the structural variants) are shown in magenta. The data are smoothed with Locally Weighted Scatterplot Smoothing (LOWESS). **C.** Distribution of SNPs along the alignment. A split into biallelic SNPs with MAF ≥ 0.01 from the alignment and recovered SNPs is shown in gray and magenta, respectively. SNP density is non-uniform across the alignment, with notable enrichment near specific mitochondrial gene regions (See **D**). **D.** SNP assignment to parental haplotypes indicates structural rearrangements and potential recombination events. The map of SNP location in the alignment. **E.** The selection of sequences and SNPs passing filtration (See Methods). The seven most abundant sequence clusters are labeled as potential parents, and the corresponding SNPs are colored according to the potential parental sequences. SNP #84 (position 27734 of the reference mitogenome) is highlighted with arrows. **F.** Genome annotation showing positions of SNPs relative to mitochondrial genes. SNP #84 corresponds to position 27734 of the reference mitogenome and is located in the *cytB* gene, which is well known to confer resistance to QoI fungicides (the gene is on the negative “Crick” DNA strand).

### Mitochondrial gene content turnover in the Zymoseptoria clade

To recapitulate mitochondrial genome evolution predating the structural variation observed within the species, we assembled genomes of four additional species of the *Zymoseptoria* genus (*Z. pseudotritici, Z. brevis, Z. ardabiliae*, and *Z. passerinii*). Despite the recent origin of the species (10,500 years for the split between *Z. tritici* and *Z. pseudotritici* ^56^ and 22,000 years for the split with *Z. ardabiliae* ^57^), mitochondrial genomes have diverged substantially (**Figure 6A**). Gene content, gene order, and genome sizes vary considerably within the genus, with a minimum and maximum size of 44 kb (in *Z. tritici*) and 62 kb (in *Z. ardabiliae*), respectively (**Figure 6C**). The large rearrangements are characterized by distinct ∼10 kb inversions in *Z. brevis* and *Z. ardabiliae* (**Figure 6A**, **C**). Multiple insertions, deletions and duplications affected both coding and non-coding regions. For instance, we found duplications of *atp9* and *nad6* in *Z. ardabiliae*, and *atp6*, *atp8* and *atp9* duplications in *Z. passerinii* (**Figure 6A**, **C**). A putative RNA polymerase gene is conserved between *Z. tritici* and *Z. pseudotritici,* but is absent in the other species (**Figure 6C**). Moreover, comparison of the first rearrangement hotspot of *Z. tritici* with the closest species *Z. pseudotritici* showed a break in synteny in this region (**Figure 6B**). There, we identified three loci with 450 bp repeats resembling the repeat structure discovered in the *Z. tritici* haplotype 3. Segments in between repeats encoded a truncated copy of *nad2,* as well as a translocation of the *atp6–atp8* gene segment (**Figure 6B**). The segments also encoded species-specific sequences such as the ORF584 (*Z. pseudotritici* only) and 795-aa COX1-containing ORF (unique the *Z. tritici* haplotype 3). The genomic context of the second hotspot was well preserved between *Z. tritici* and *Z. pseudotritici* mitogenomes, but haplotype 4 insertion is absent outside *Z. tritici*. Notably, structural variation in the *Z. pseudotritici* mitogenome also affects NAD2-encoding regions. This suggests that the *nad2* duplication may be the ancestral state, rather than the configuration found in *Z. tritici* haplotype 1, which lacks has only one copy. These findings confirm that the *Zymoseptoria* mitogenomes have persistent genomic fragility in specific regions underpinning both within-species rearrangements and between-species synteny breaks.

**Figure 6.**
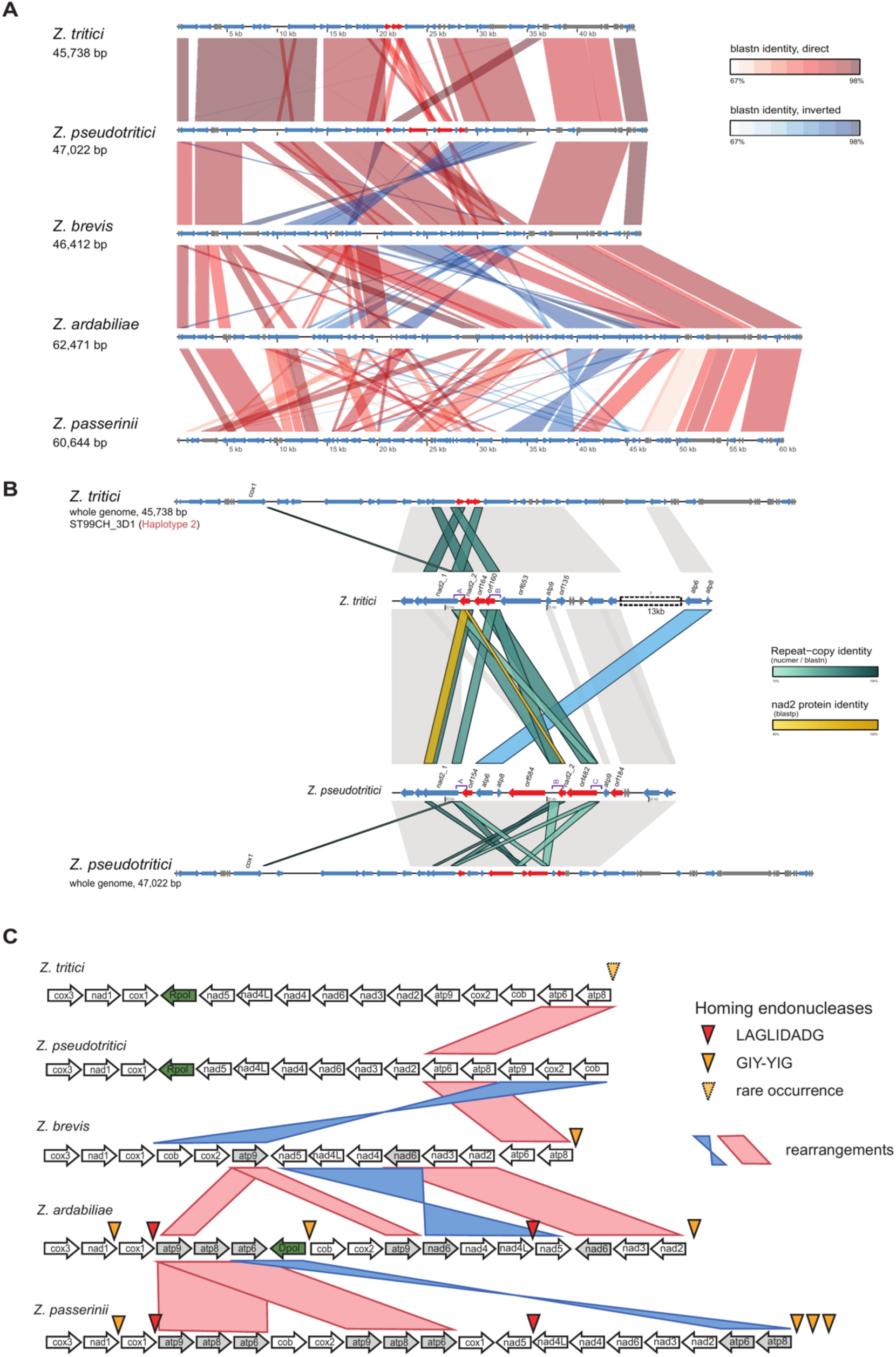
Mitogenome diversity across the *Zymoseptoria* genus, evolution of mitochondrial selfish elements between species. **A.** Synteny plot for mitochondrial genomes of *Z. tritici* (reference genome, isolate IPO323) and 4 other species in the *Zymoseptoria* genus, with the mitogenome length. All annotated protein-coding genes are depicted in blue, tRNA and rRNA genes in grey. The genes of the indel in the first rearrangement hotspot of *Z. tritici*, as well as the genes from the same locus in *Z. pseudotritici,* are depicted in red. Direct synteny blocks are colored in shades of red depending on the identity level, and inversions are colored in shades of blue. The dramatic genome size variation and the rapid breakdown of synteny can be observed. **B.** Synteny of the first rearrangement hotspot between the *Zymoseptoria tritici* haplotype 2 (ST99CH_3D1) and *Z. pseudotritici*. From top to bottom, the tracks show the complete ST99CH_3D1 mitochondrial genome (45,738 bp); an enlarged view of the ST99CH_3D1 hotspot-1 region, which carries the 1,171 bp short-allele insertion; the corresponding region of *Z. pseudotritici*; and the complete *Z. pseudotritici* mitochondrial genome (47,022 bp). Arrows show annotated genes; coloring is the same as in panel A. Grey blocks show main collinear regions identified by blastn. Teal ribbons show nucleotide identity between repeated and paralogous sequences (blastn, 70–100%), including the two 603 bp direct repeats (1 and 2) that flank the insertion in ST99CH_3D1. Gold ribbons show protein identity from blastp (80–100%). ST99CH_3D1 nad2_copy2 is 87.3% identical to the *Z. pseudotritici* nad2_copy2 fragment. The blue ribbon shows the translocation of the atp6/atp8 genes in *Z. pseudotritici*. Alignments shorter than 200 bp are not shown. **C.** Gene order graph for five species in the *Zymoseptoria* genus. Only essential protein-coding genes are depicted, with parts of the syntenic blocks. Duplicated genes are colored in grey. DNA and RNA polymerase genes are depicted in green. The red and yellow arrows represent the presence of homing endonuclease genes from the LAGLIDADG and GIY-YIG families, respectively.

### Evolutionary dynamics of selfish elements

Selfish genetic elements were found in the majority of analyzed mitogenomes (**Figure 6B**, **Supplementary Figure 8**), including several LAGLIDADG homing endonucleases (two copies each in *Z. ardabiliae* and *Z. passerinii*). One copy in *Z. ardabiliae* is flanked by group I introns. One GIY-YIG homing endonuclease was found in *Z. brevis*, three in *Z. ardabiliae,* and four in *Z. passerinii*. The insertion point of GIY-YIG is conserved across species carrying the gene (**Supplementary Figure 8)**, though, unlike the intact *Z. tritici* GIY-YIG ORF (233 aa), the orthologous copies in the sister species are degraded. The *Z. brevis* copy has a single N-terminal frameshift, the *Z. ardabiliae* copy has two frameshifts and a premature stop codon that cuts off its C-terminus, and the *Z. passerinii* copy is a pseudogene split across all three reading frames with an internal stop codon. *Z. passerinii* carries an additional GIY-YIG gene at a 691 bp distance. However, the sequence shares only weak similarity to the GIY-YIG copy shared with *Z. tritici* **(Supplementary Figure 8**). The rarity of the GIY-YIG copy in *Z. tritici* (0.4% globally and restricted to Eastern Europe), as well as the likely absence in the closest sister species *Z. pseudotritici*, suggests a horizontal transfer event or strong selection for GIY-YIG loss in the clade leading to *Z. tritici*.

To investigate the evolutionary origins of the *Z. tritici* GIY-YIG, we searched for protein sequence homology in databases. Furthermore, we annotated all available mitogenomes of Ascomycota for ORFs encoding the GIY-YIG domain. We clustered the retrieved GIY-YIG homologs based on the pairwise alignments and constructed a phylogenetic tree of the closest homologs to the *Z. tritici* copy (**Figure 7B**). Beyond *Zymoseptoria* mitogenomes, we found highly similar GIY-YIG in the *Cladosporium* and *Pseudocercospora* genera of the Dothideomycetes. Further homologs were found in Lecanoromycetes, Sordariomycetes, and Orbiliomycetes. To assess signatures of horizontal transfer events, we constructed a species tree based on concatenated BUSCO genes (**Figure 7A**). GIY-YIG copies from the genus *Zymoseptoria* were clustered with GIY-YIG homologs of *C. sphaerospermum* rather than members of the closely related *Pseudocercospora* genus. Within Dothideomycetes, GIY-YIG homologs split into two well-supported clades, one of which grouping with homologs from Sordariomycetes and Orbiliomycetes. The phylogenetic patterns of GIY-YIG homologs in Ascomycota may be explained by the widespread loss of multi-family GIY-YIG copies, leaving only a few clades with extant copies. However, the rarity of the *Z. tritici* GIY-YIG homolog among distantly related clades may be more parsimoniously explained by horizontal gene transfer events.

**Figure 7.**
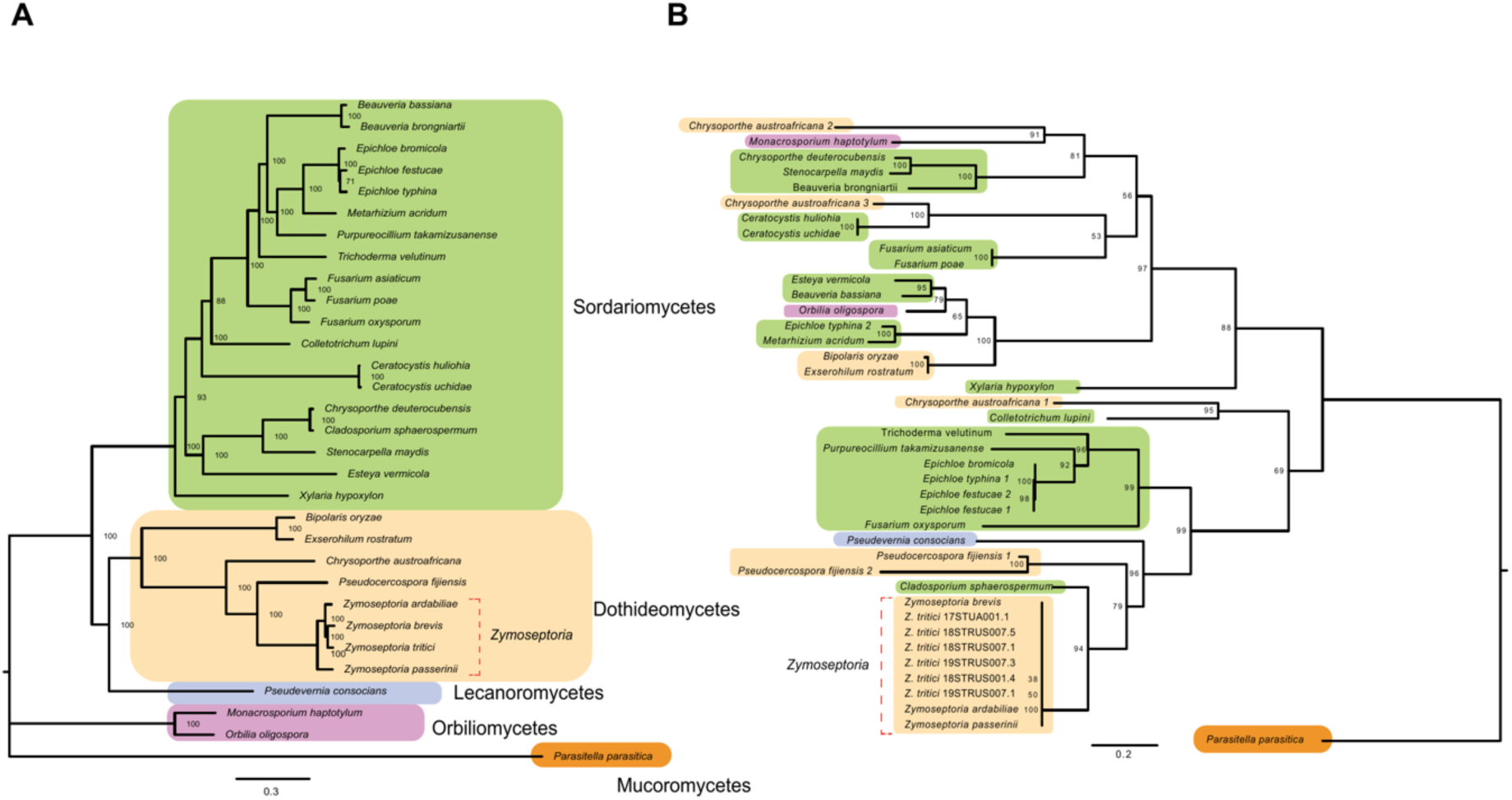
The evolutionary origins of the *Z. tritici* GIY-YIG. **A.** Maximum likelihood tree for the species with the closest homologs to the GIY-YIG homing endonucleases found in *Z. tritici*. The tree is based on the alignment of 100 BUSCO genes. Species from the Zymoseptoria genus are marked by a red bracket. The species from the class Dothideomycetes are colored in light orange, from Lecanoromycetes in blue, from Sordariomycetes in light green, and from Orbiliomycetes in purple. *Parasitella parasitica* from the class Mucoromycetes was used as the outgroup and colored in orange. **B.** Maximum likelihood tree of GIY-YIG homing endonucleases, based on amino acid sequences from 6 isolates of the *Z. tritici* population; from *de novo* assembled mitochondrial genomes of *Z. brevis, Z. ardabiliae,* and *Z. passerinii* (*Zymoseptoria* genus, yellow); and closest homologs from other species. The coloring scheme for the isolates repeats the coloring by classes on the species tree. If more than one copy of GIY-YIG per species was included, the copies are distinguished by a number after a species name.

## Discussion

Here, we focused on analyzing mitogenome variation generated over short evolutionary time spans within a species and close relatives. The deep genomic survey revealed nuclear-mitochondrial discordance, evidence for recombination, and the presence of an orphan, geographically restricted GIY-YIG selfish element. Tracing the origin of this element revealed likely horizontal gene transfer events among distantly related fungi.We established a comprehensive map of mitochondrial genome variability based on over 2000 genome sequencing datasets. Structural variants accounted for 19% (**Supplementary Figure 6B**) of the mitochondrial genome, with genome size ranging from 43.9 to 52.3 kb. This is lower than in other well-studied fungal species, such as *F. oxysporum* and *S. cerevisiae,* with variants accounting for 87% (genome size 37-69 kb) and 32% (73-96 kb) of genome size variation, respectively. Between species, differences in effective population size (*N*ₑ), life history, and reproductive strategies are expected to influence the observed mitochondrial genetic diversity ^58,59^. Species with a larger Nₑ should maintain higher mitochondrial variation due to weaker genetic drift. Reproductive strategies affect the opportunity for mitochondrial heteroplasmy during mating and, therefore, also affect the likelihood of recombination among mitochondrial genomes. The factors that drive the occurrence of heteroplasmy are unknown, and in most species mitochondrial inheritance is strictly maternal ^60^. Similarly, mechanisms facilitating mitochondrial fusions and the occurrence of recombination remain poorly understood. The three species differ in these traits. *Z. tritici* undergoes at least one sexual cycle per year and is thought to be highly diverse based on nuclear linkage disequilibrium decay and polymorphism analyses ^61^ ^62^. The species is also young, with a divergence time estimated to be 10,500 years^56^. In contrast, *F. oxysporum* lineages are far more divergent across the species complex, with a divergence time estimated at ∼500,000 years and a high SNP density on core chromosomes (57 SNPs/kb) ^63^. However, the effective population size (*N*ₑ) is difficult to assess for *F. oxysporum* given its predominantly clonal reproduction. Species with a dominant clonal population structure, such as *F. oxysporum* and *S. cerevisiae,* can complicate direct comparisons of nuclear and mitochondrial diversity. In contrast, regularly outcrossing species and accessible sampling of individual populations, such as for *Z. tritici,* enable a more direct assessment of evolutionary forces acting upon mitochondrial polymorphism.

Mito-nuclear incongruence is a puzzling feature of mitochondrial genome evolution at the species level and has been described in the best-sampled fungi to date, such as *F. oxysporum* and *S. cerevisiae.* Here, we show that incongruence has also arisen in a sexually reproducing species with a well-characterized evolutionary history. The nuclear genome of *Z. tritici* reflects the colonization history tied to the spread of wheat cultivation: from the Middle East to Europe and North Africa, and subsequently across the globe. In contrast, mitochondrial population structure is largely discordant based on both PCAs and sNMF clustering. The discordance may have arisen already in the Middle Eastern populations if these harbored all major haplotypes (V1-3). These haplotypes were then likely spread to North Africa and persisted in both regions. A colonization bottleneck may have led to only one mitochondrial haplotype (V1) successfully colonizing Europe and subsequently dispersing to North America and Australia. Meanwhile, another mitochondrial haplotype (V3) established itself on the Iberian Peninsula, either through a separate migration event from North Africa or directly from the Middle East. This V3 haplotype may have later spread to Central and South America, where it became predominant. Present-day Eastern Europe represents a contact zone where V1 and V2 mitochondrial haplotypes coexist. If bottlenecks were the primary driver of mitochondrial haplotype differentiation, the discordance with patterns at nuclear loci may have arisen from the stronger bottlenecks imposed on the uniparentally inherited mitochondria. However, selection favoring specific haplotypes during colonization events may have also contributed to the observed incongruence. Present-day selection is best understood for the emergence of QoI fungicide resistance, but multiple mitochondrial haplotypes share the same resistance mutation. Historic selection pressures may nevertheless have favored specific mitochondrial haplotypes.

Mitochondrial genome variation, such as that observed in *Z. tritici* differentiation into major haplotypes, arose predominantly from variation in intergenic regions and chromosomal inversions. In *F. oxysporum*, *de novo* intron insertions and long variable regions (LVRs) were the primary sources of structural variability, whereas in *S. cerevisiae* chromosomal inversions were additional contributors to structural variation ^64^. The structural variants we observed in *Z. tritici* are dominated by two rearrangement hotspots. In particular, the flanking repeats and truncated *nad2* copies in Haplotypes 2 and 3 suggest that the first hotspot arose through tandem duplication–random loss (TDRL). Under this model, a segment of the mitogenome is duplicated in tandem, after which redundant gene copies are progressively lost or degenerate into non-functional remnants^65,66^. TDRL is well established as a major driver of gene order change in metazoan mitogenomes, where rearrangement hotspots and residual pseudogenes are its hallmark signatures ^67^, and it can generate distinct gene orders even within a single species^68^. In fungi, structural rearrangements have mainly been attributed to non-homologous recombination^69,70^ and to the activity of mobile elements ^71^. While *Z. tritici* mitochondria are essentially devoid of selfish elements, our finding that recombination is concentrated in hotspots that co-localize with structural rearrangements suggests that recombination itself generates at least part of the mitogenome variation. Because the first hotspot is flanked by repeats, repeat-mediated recombination likely facilitated the initial duplication. TDRL and recombination are therefore complementary rather than mutually exclusive explanations.

The paucity of selfish elements in *Z. tritici* mitochondria was striking compared to species with more structurally diverse mitochondrial genomes. We identified only six isolates (0.4%), all originating from Eastern Europe, encoding an HE and explaining 5% of the observed genome size variation. We retraced the origin of this GIY-YIG element to close sister species. The closest homologs present in sister species and the largely conserved insertion sites are consistent with a model of GIY-YIG presence being ancestral to the investigated species of the *Zymoseptoria* genus. In some species, the GIY-YIG element multiplied and generated additional copies (*Z. ardabiliae* and *Z. passerinii*), whereas in others it was likely lost (*Z. pseudotritici*) or retained only in a small fraction (*Z. tritici*). Consistent with widespread gene loss, the nearest homolog to the *Zymoseptoria* GIY-YIG copy was found in *Cladosporium sphaerospermum*, implying broad losses in more closely related species. Some GIY-YIG copies found in Dothideomycetes clustered with GIY-YIG from either Sordariomycetes or Orbiliomycetes. These copies represent distinct GIY-YIG paralogs, which underwent widespread losses in Dothideomycetes. Conversely, we found GIY-YIG copies in species from the classes Lecanoromycetes and Sordariomycetes on well-supported branches clustering with the *Zymoseptoria* clade. Taken together, we observed clear incongruence between the phylogenies of GIY-YIG sequences and nuclear loci. Beyond strong evidence for paralog formation and multiple losses, horizontal transfers likely contributed to the observed distribution, as neither paralog formation nor widespread GIY-YIG gene losses parsimoniously explain the mismatch in GIY-YIG and species phylogenies. The clearest signal of horizontal gene transfer comes from the GIY-YIG copies found in the genus *Pseudocercospora* clustering with *C. sphaerospermum* even though *Pseudocercospora* is phylogenetically closer to *Zymoseptoria*. Furthermore, the clustering of GIY-YIG copies from distantly related classes (*i.e.,* Lecanoromycetes) provides additional support for a horizontal transfer event.

The evolution of selfish elements in mitochondria reveals complex patterns, characterized by frequent duplications across species and high divergence among mitochondrial haplotypes. Possible explanations for these patterns include the likely low selection pressure on selfish element proliferation, the presence of defense mechanisms limiting their propagation, or selection acting on mitochondrial genome size. Among such mechanisms, nuclear-encoded defense systems against transposable elements, such as RIP ^72^ and piRNA ^73^, are not known to be functional on repeats in mitochondrial genomes. Counteracting forces against selfish element proliferation beyond the costs of genome expansion remain unknown. However, the predominantly uniparental inheritance of mitochondria may itself serve as a defense mechanism against selfish element proliferation by protecting distinct mitochondrial haplotypes from exposure to each other ^74^. Selfish elements may also spread by horizontal transfer between species, through mechanisms that remain unclear but may involve selfish mitochondria ^75^ and cell-to-cell anastomosis ^76^. Although there are numerous examples of horizontal gene transfer (HGT) of essential mitochondrial genes from mitochondrial to nuclear genomes ^77^, the evidence for the transfer of mitochondrial selfish elements remains scarce. Only isolated cases of selfish element transfer to the nuclear genome have been described ^78^. To date, several studies have provided indirect evidence for horizontal transfer of mitochondrial selfish elements in fungi, based on phylogenetic incongruence between selfish elements and host species trees, particularly for GIY-YIG and LAGLIDADG homing endonucleases ^79–83^. Therefore, the extensive horizontal transfer observed for nuclear transposable elements ^84,85^ may also extend to mitochondrial selfish elements. Our analysis similarly examined the evolutionary history of a specific family of GIY-YIG endonucleases and revealed multiple putative HGT events across the fungal phylogeny. However, these inferences may be influenced by the highly patchy distribution and sampling of selfish elements. Large-scale population and comparative genomics analyses are key to unraveling mitochondrial evolution at even finer resolution.

## Methods

### Isolate collection, culturing, and sequencing

We analyzed the following whole-genome *Z. tritici* sequencing datasets: 1106 worldwide isolates sequenced by Illumina ^47^; 1087 European isolates sequenced by Illumina ^48^; 23 isolates sequenced by PacBio, from ^86^ and BioProject PRJNA593239. Furthermore, we analyzed PacBio sequencing data of four sister species including *Z. pseudotritici, Z. brevis, Z. ardabiliae* and *Z. passerinii* ^87^. Information about all geographic origins and collection dates is summarized in Supplementary Table 1. The data were originally generated by culturing the fungal isolates on various media, including V8, yeast sucrose broth (YSB), yeast malt agar (YMA), or in liquid culture before DNA extraction (see references above). DNA extraction was performed using QIAGEN kits, and sequencing libraries were prepared using TruSeq library preparations for the Illumina platform. For PacBio sequencing of *Z. tritici* isolates and sister species, detailed procedures are described in ^86^. In brief, lyophilized spores were used to extract high molecular weight DNA following a modified CTAB protocol developed for plant tissue. PacBio SMRTbell libraries were prepared from 15-30 μg of DNA.

### Variant calling

Raw Illumina reads of each of the 2193 isolates were trimmed by Trimmomatic v.0.39 with default settings ^88^ and mapped against the IPO323 reference genome with bowtie2 v.2.4.1 ^89^, following the steps described in ^47^. Mapped read files were coordinate-sorted with samtools v.1.19.2 ^90^; variant calling, variant selection, and filtration were performed using the Genome Analysis Toolkit (GATK) version 4.4.0.0^91^. Variants were filtered by vcftools v.0.1.16 ^92^, following the steps described in ^47^. For both the nuclear and mitochondrial datasets, only biallelic SNPs were retained. Variants were filtered to allow a maximum of 80% missing data per site and 1% missing data per sample, with a minimum minor allele frequency (MAF) of 0.02. Nuclear SNPs were further thinned to one SNP per 1,000 bp to reduce linkage effects.

### Population genetic structure

To filter SNPs for population structure analysis, we used vcftools v.0.1.16 ^92^. The SNP set obtained for the nuclear genome comprised 13,326 biallelic SNPs with a minor allele frequency of 2% and was subsampled to 1 SNP per 1000 bp (--thin 1000). Principal component analysis (PCA) was performed using the SNPRelate package ^93^. The sNMF clustering method from the LEA R package ^94^ was used. For the nuclear genome, the clustering analysis was conducted for 1-15 K values (number of clusters) with 10 repetitions per K. To determine the optimal K, we employed the entropy method implemented in sNMF, which assesses the model’s fit to the data, the smallest cluster size, and the number of non-admixed isolates assigned to any cluster with a coefficient exceeding 0.75. For the mitochondrial genome, we analyzed 1237 SNPs for the same dataset of 2193 isolates. Out of all SNPs, 378 were biallelic and genotyped in all isolates. Applying an additional minor allele frequency filter of 2% reduced the number of SNPs to 17. This subset of 17 SNPs was used for both sNMF-clustering and PCA. Switching between the larger SNP set of 378 loci and the small one (*n* = 17 SNPs) did not meaningfully affect the PCA (**Supplementary Figure 1 G, H**).

### Mitochondrial genome assembly

To assemble mitochondrial genomes for the 23 *Z. tritici* isolates sequenced with PacBio (**Supplementary Table 2)**, as well as four isolates of other *Zymoseptoria* species (**Supplementary Table 4)**, we conducted a two-step approach: we first filtered all raw long reads by mapping against the IPO323 reference mitochondrial genome ^50^ using minimap2 v.2.26-r1175 with default parameters ^95^. Using only mapped reads, we used Canu v.2.2 ^96^ to assemble mitochondrial genomes. Canu settings included an estimated genome size of 0.1 Mb (-genomeSize=0.1m), a filter for minimal read length of 500 bp (-minReadLength=500), and a maximum allowed error rate in the corrected reads of 0.045 (-correctedErrorRate=0.045). If no single circular contig was obtained, we attempted to assemble the mitochondrial genome using Flye v.2.9.2-b1786 ^97^ with an estimated genome size of 0.1 Mb (--genome-size 0.1m). Successfully assembled mitochondrial genomes were reverse complemented and adjusted to the IPO323 reference assembly if needed.

We assembled additional mitochondrial genomes using paired-end Illumina reads of 2,193 isolates (**Supplementary Table 1)**, by GetOrganelle v1.7.7.0 ^98^ based on the nucleotide sequence of the *cox1* gene of the IPO323 mitochondrial genome as the seed, the *fungus_mt* option as the reference database, maximum extension rounds (-R) of 10, and a range of k-mers (-k) of 21, 45, 65, 85 and 105. To align and compare mitogenomes, we reverse-complemented and position-adjusted contigs according to the IPO323 reference. A total of 2061 isolates were successfully assembled.

### PCR validation of the insertion

The presence of the insertion in the ST95CH_3B2a strain and the absence of it in IPO323 were validated by PCR using two independent primer pairs designed to amplify the insertion site. Primer pair 1 (1_F: 5′-CTAAGTCCGCTGTGCTTA-3′ and 1_R: 5′-ACCTGCCTGAACTACCTG-3′) generated an expected amplicon of 897 bp in the reference allele and 2,671 bp in the insertion-containing allele. Primer pair 2 (2_F: 5′-GGAACGGTGCTGCACTTAC-3′ and 2_R: 5′-GCAAATCCGAGGAGATCG-3′) generated an expected amplicon of 1,428 bp in the reference allele and 3,202 bp in the insertion-containing allele. PCR amplifications were performed using GoTaq G2 DNA Polymerase. The PCR program consisted of an initial denaturation step at 95°C for 2 min, followed by 30 cycles of denaturation at 95°C for 30 s, annealing at 52°C for 30 s, and extension at 72°C for 4 min 30 s. A final extension step was performed at 72°C for 5 min, followed by a hold at 4°C. PCR products were separated on a 0.8% agarose gel by electrophoresis at 75 V for 1 h. The presence or absence of the insertion was determined based on the observed amplicon sizes.

### Annotation of mitogenomes

To annotate *Z. tritici* mitogenome assemblies, we utilized Mitos2 v.2.1.0 ^99^ with the genetic code set to 4 (-c 4) and fungal proteins and RNA sequences as the reference database (-r). To validate protein predictions, we performed hmmsearch (HMMER 3.2.1) ^100^ with the Pfam hmm-database for all predicted open reading frames, with an e-value threshold (-E) set to 1e-10. Following this approach, we reannotated the reference mitochondrial genome of *Z. tritici* (the isolate IPO323) and identified a previously undescribed gene with an RNA-polymerase protein domain, located between cox1 and nad5 genes. To annotate variable regions among *Z. tritici* genomes, we searched for additional ORFs using ORFfinder v.0.4.3 ^101^ (genetic code 4) and scanned all predicted ORFs for known protein domains with InterProScan ^102^. For the sister species mitogenome assemblies, we additionally used the online version of the annotation tool MFannot ^103^ with the genetic code set to 4. To convert .sqn files to GenBank .gb files, we used the tool asn2gb ^104^. To screen for homing endonucleases, we performed blastn ^105^ searches using the *Z. tritici* GIY-YIG gene as the query.

### Construction of the pangenome graph and structural variant detection

To construct a mitochondrial pangenome graph, we first excluded assemblies with genome sizes above 55 kb (*n =* 2 assemblies) and below 42 kb (*n =* 86 assemblies). For the remaining assemblies, we applied a similarity filter based on a multiple sequence alignment performed with mafft ^106^. We excluded sequences generating more than 4000 gap columns with a gap frequency greater than 0.01, which resulted in the removal of 19 assemblies. The filtered dataset of 1977 assemblies (1954 isolates, sequenced by Illumina, and 23 isolates, sequenced by PacBio) was then used to construct the pangenome graph.

We used the pangenome graph builder (PGGB) version 0.6.0 ^107^ with a 95% identity setting for mapping and alignment (-p 95). The graph was visualized using Bandage version 0.8.1 ^108^. We used odgi ^109^ v.0.8.2 to obtain the depth of each node (odgi depth). To annotate the graph in Bandage, we performed BLAST searches of the genes annotated in the 1977 *Z. tritici* mitochondrial genomes. To evaluate variants in the graph and generate a VCF file, we used the *deconstruct* command from vg-tools version 1.65.0 ^110,111^ with IPO323 as the reference, specifying the flag -n to obtain nested variants and the flag -L to cluster together the alleles with 80% similarity (-L 0.8). To report structural variants, we applied a filter of 30 bp for the difference between the reference and any alternative alleles, as well as between alternative alleles. The mitochondrial haplotypes were defined based on unique allele combinations across SNP and SV loci (defined as “unique” haplotypes in Supplementary Table 1) or only based on SV loci (termed “SV haplotypes”).

### Recombination detection

We aligned the 1977 mitochondrial genome sequences using MAFFT and default settings (-- auto). To avoid inflating the alignment, sequences creating gap columns for more than 1000 total positions with a gap frequency of less than 0.01 were removed. The resulting alignment was re-aligned with MAFFT using --auto and trimmed with trimAl 1.5.0 using the -strict option. Trimmed alignment columns were masked with gaps to retain the original distance in the alignment (gap-distance-correction). The alignment was converted to VCF using SNP-sites ^112^ (snp-sites -mvp). SNP positions (*n*=131) and haplotypes were retrieved from the VCF file using Phylip format files generated by SNP-sites. VCFtools ^92^ was used for linkage disequilibrium analyses of biallelic SNPs with MAF ≥ 0.01 (*n*=111) (--maf 0.01 --max-alleles 2 --min-alleles 2 --hap-r2). Linkage disequilibrium r^2^ decay was plotted using circular genome coordinates with Locally Weighted Scatterplot Smoothing (LOWESS; frac = 0.25). VCFtools was used to extract the same biallelic SNPs used for linkage disequilibrium analysis (--maf 0.01 --max-alleles 2 --min-alleles 2 --kept-sites).

To expand the set of available biallelic SNPs, we analyzed the above-described sequence alignment before masking, and added 20 SNPs from structural variants absent in the reference genome. The alignment was filtered to remove (first) sequences and (second) positions exceeding a threshold of 10 gaps, followed by pairwise Hamming distance calculation between all sequences, ignoring gap positions. Ward’s hierarchical clustering with a distance threshold of 0.1 was used to identify major parental clusters. From these clusters, consensus sequences were generated by selecting the most common nucleotide at each position. Each position in every sequence was then assigned a parental identity based on nucleotide matching to parent consensus sequences using a priority-based claiming system, where the largest cluster (Parent 1) had the highest priority. Sequences not belonging to parental clusters were classified as putative recombinants. This full-length sequence analysis was followed by filtering steps that removed invariant positions, highly similar sequences (99% identity threshold), and retained only biallelic SNPs containing exactly two different parental signatures.

We used the recombination detection tools RDP, GENECONV, BOOTSCAN, MAXCHI, CHIMAERA, SISCAN, PHYLPRO, LARD, and 3SEQ implemented in the RDP5 program^113^ with default settings on the preprocessed alignment (filtered, masked, and gap-distance-corrected alignment blocks). ARGweaver ^114^, fastGEAR ^115^, and PhiPack ^116^ with default settings were used to further assess signatures of recombination. SplitsTree version 6.0.0 ^117^ was used to perform a phylogenetic network analysis.

### Phylogenetic analyses

To identify the closest homologs of the *Z. tritici* GIY-YIG homing endonuclease, we pursued two approaches: (1) we performed a blastp ^105^ search and filtered hits for query coverage >69% and identity >35% resulting in 10 retained hits; (2) we downloaded all available Ascomycota mitogenomes from NCBI (*n =* 709) and annotated these using MITOS2 v.2.1.0 ^99^, then assessed the domain architecture of all predicted ORFs based on hmmsearch (HMMER 3.2.1) ^100^, resulting in 686 ORFs encoding GIY-YIG domains. Using pairwise Smith-Waterman alignments, we clustered GIY-YIG amino acid sequences resulting from the blastp search of the sister species in the *Zymoseptoria* genus and resulting from the analysis of all available Ascomycota mitochondria.

To contrast the GIY-YIG phylogeny with the species phylogeny, we filtered the top 50 most similar homologs for the availability of a corresponding nuclear genome assembly. This resulted in retaining 46 amino acid sequences from 31 species (Supplementary Table 5). Then, we aligned GIY-YIG protein sequences using MAFFT v7.520 ^106^, trimmed the alignment using trimAl v1.4.rev15 ^118^, and constructed a phylogeny using IQtree2 v.2.2.6 ^119^. We performed 1000 bootstrap replicates (-bb 1000). ModelFinder identified the VT+F+I+G4 substitution model as most appropriate. To generate a species phylogeny, we first performed BUSCO analyses of the nuclear genome assemblies and then randomly selected 100 BUSCO genes. We aligned BUSCO genes independently using MAFFT v7.520 ^106^, then concatenated the alignments using PhyKIT v2.1.2 ^120^, trimmed the alignment using ClipKit v.2.7.0 ^121^, and constructed the maximum likelihood tree by IQTree2 v.2.2.6 ^119^ with 1000 bootstrap replicates (-bb 1000) and *Parasitella parasitica* (Mucoromycetes) as the outgroup (-o). ModelFinder identified the Q.yeast+F+I+R6 substitution model.

## Declarations

### Data Availability

To analyze mitogenomes, we used the following whole-genome Z. tritici sequencing datasets: 1106 worldwide isolates sequenced by Illumina ^47^; 1087 European isolates sequenced by Illumina ^48^; 23 isolates sequenced by PacBio, from ^86^ and BioProject PRJNA593239. Mitochondrial genome assemblies are available on GenBank: accessions PZ827084-PZ827109 for PacBio-sequenced genomes, PZ825129-PZ827082 for Illumina-sequenced genomes. The genome annotations are published on Zenodo with DOI: 10.5281/zenodo.22097545. All other data can be found in the Supplementary Information or referenced in the Methods section.

## Acknowledgments

We are grateful to Mélanie Echernier for assistance with resistance analyses, Anouk C. van Westerhoven for sharing scripts for the pangenome analysis, Tobias Baril and Thomas Wicker for advice on phylogenetic analyses, and Christina Lado Arcos and Raissa Da Silva Almeida for help with laboratory work.

## Author contributions

I.S. and D.C. conceived the study; I.S., P.K., O.G., D.K. and A.F. performed analyses; A.F. and G.P. provided datasets; D.C. supervised the work; I.S. and D.C. wrote the manuscript with input from P.K. and D.K.

