## Supplementary Figures for "Drivers of intraspecies mitochondrial genome evolution revealed by deep population sequencing of a major fungal wheat pathogen"

Skakov et al.

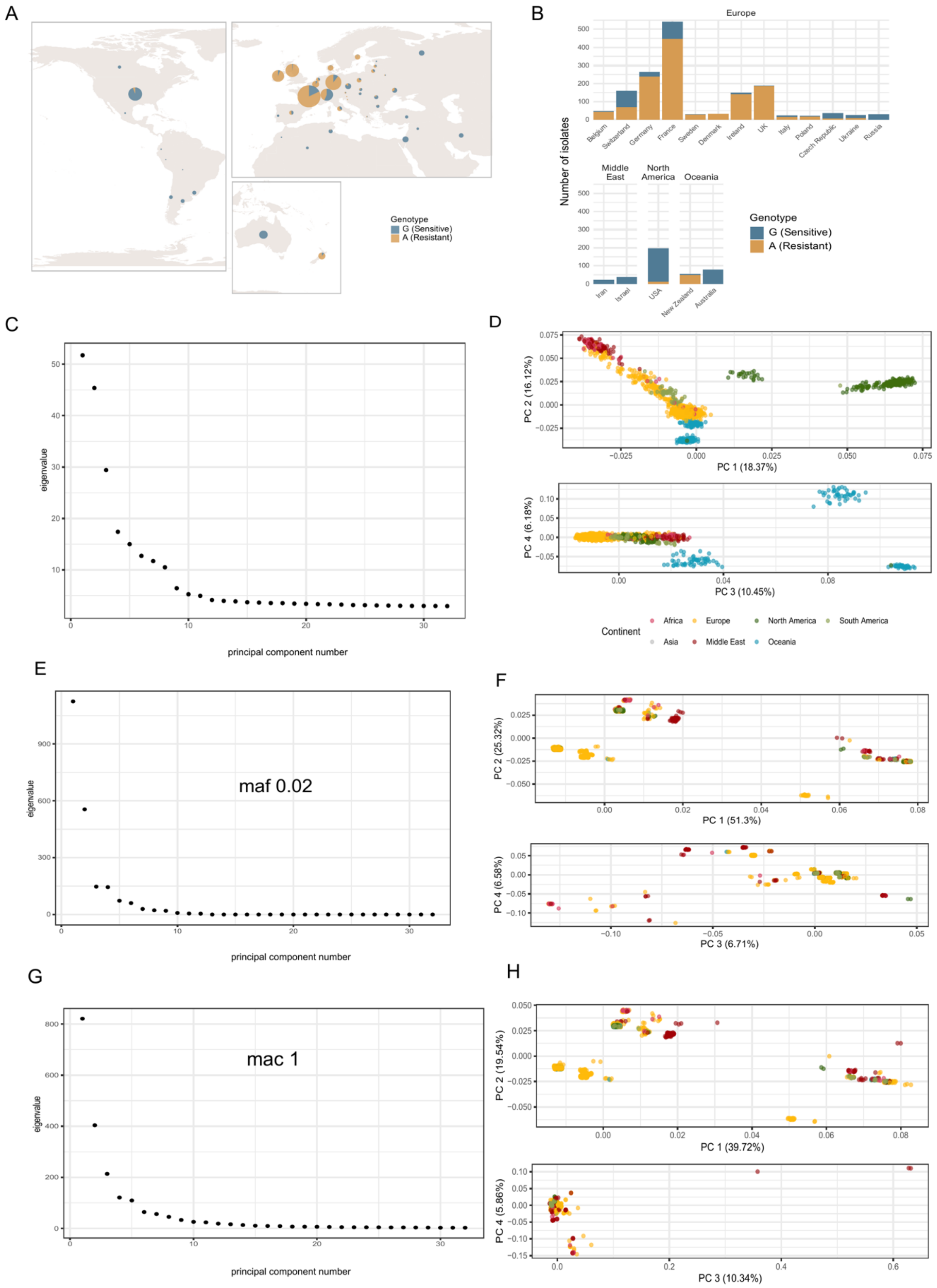


**Supplementary Figure 1.** Isolate distribution and mito-nuclear population structure incongruence by PCA. **A.** Map of the distribution among the continents and countries of the G143A mutation in the *cytB* gene, conferring resistance to QoI fungicides. The G143A mutation is predominant in Europe and New

Zealand, and absent or nearly absent in the Middle East, North America, and Australia. **B.** Absolute number of isolates with and without the G143A mutation per country. **C.** Eigenvalue distribution for each principal component for PCA on the SNPs from the nuclear genome of *Z. tritici*. **D.** Principal component analysis (PCA), showing the first, second, third, and fourth components (PCs) based on variants from the nuclear genome. Colors indicate the geographical origin of the isolates. **E.** Eigenvalue distribution for each principal component for the mitochondrial genome of *Z. tritici,* based on 17 biallelic SNPs with minor allele frequency above 0.02. **F.** Principal component analysis (PCA), showing the first, second, third, and fourth components (PCs) based on 17 biallelic SNPs from the mitochondrial genome with minor allele frequency above 0.02. Colors indicate the geographical origin of the isolates. **G.** Eigenvalue distribution for each principal component for the mitochondrial genome of *Z. tritici,* based on 378 biallelic SNPs with minor allele count above 1. **H.** Principal component analysis (PCA), showing the first, second, third, and fourth components (PCs) based on 378 biallelic SNPs from the mitochondrial genome with minor allele count above 1. Colors indicate the geographical origin of the isolates.

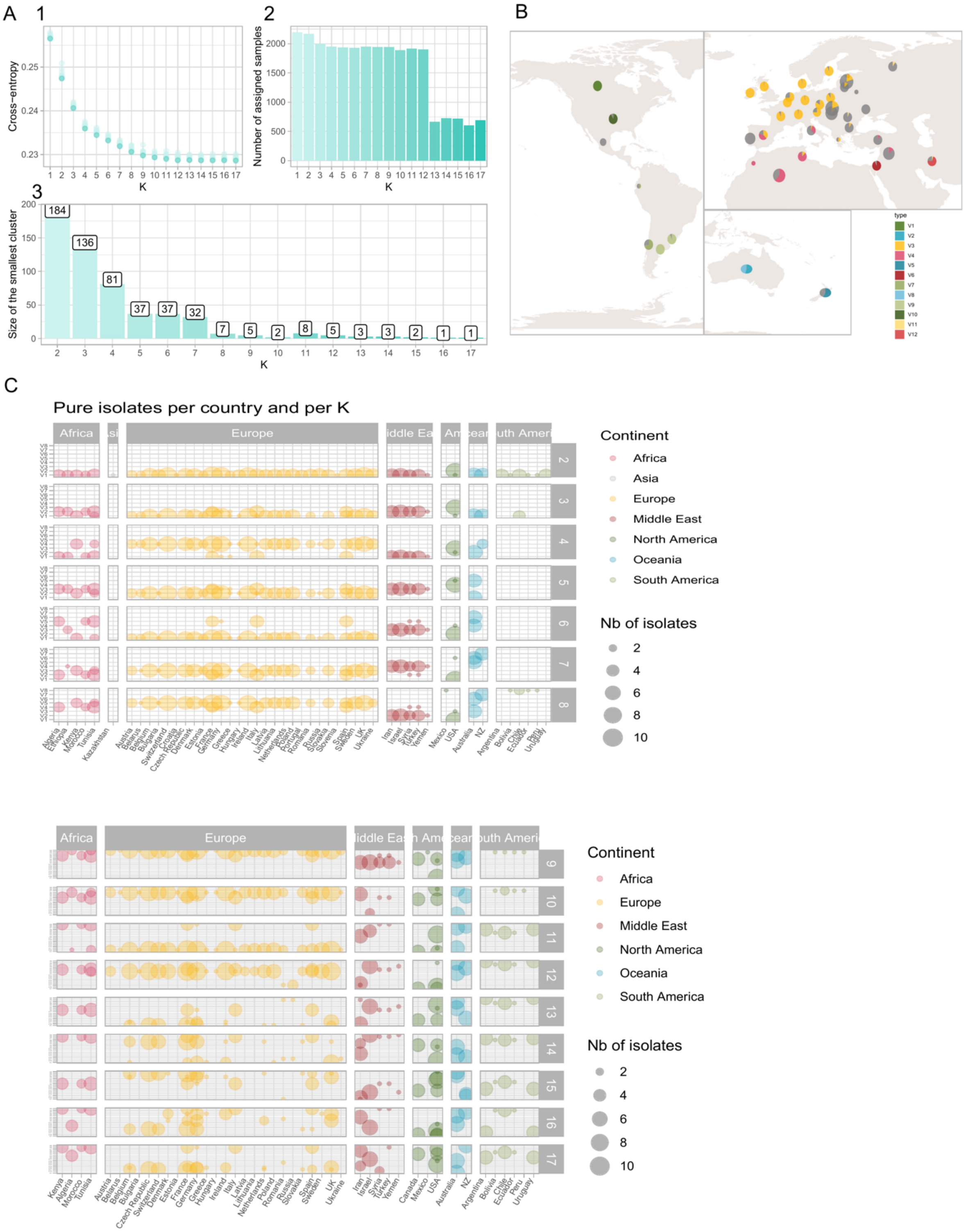

A
1
2
3
B
C
Cross−entropy
K
Number of assigned samples
Size of the smallest cluster
184
136
81
37
37
32
7
5
2
8
5
3
3
2
1
1
type
Pure isolates per country and per K
Continent
Africa
Asia
Europe
Middle East
North America
Oceania
South America
Nb of isolates
2
4
6
8
10

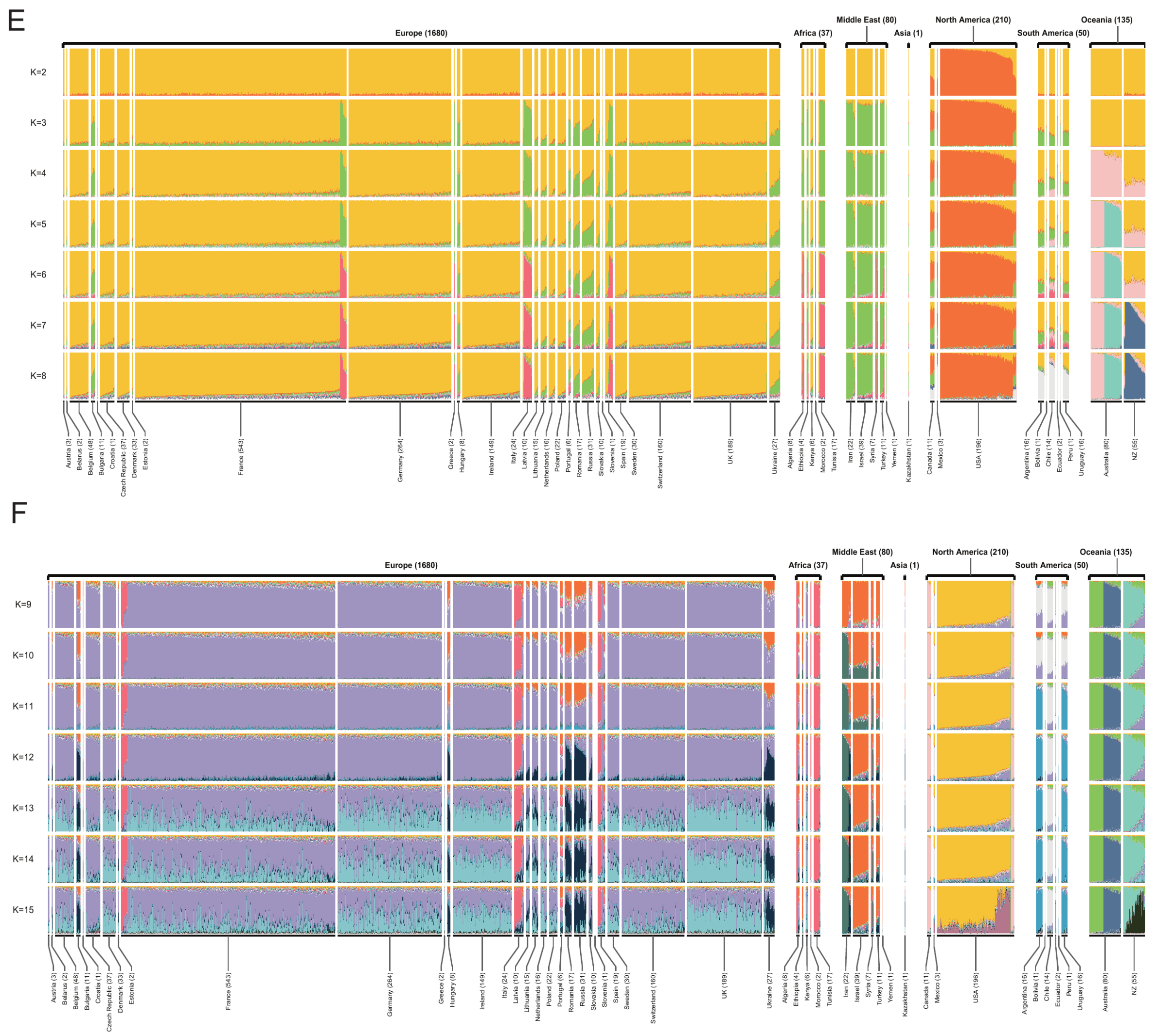


**Supplementary Figure 2**. sNMF clustering for the nuclear genomes. **A. 1.** Cross-entropy for each number of clusters (K) as assessed by the R package *LEA*. **2.** Number of isolates uniquely assigned to one cluster per K. **3.** Number of isolates assigned to the smallest cluster. **B.** Map of the genetic clustering based on a nuclear genome SNP dataset using sNMF. Each color represents a different genetic cluster, and the sizes of the slices represent the average attribution to the cluster across the isolates from each location. The isolates defined as admixed are colored in grey. Distribution of the isolates per country per cluster for the number of clusters (K) from 2 to 8 in **C** and from 9 to 17 in **D.** Genetic clustering of isolates based on nuclear SNPs showing per-isolate barplots with cluster assignments in **E, F.** Barplots for the best replicate for each K (as determined by cross-entropy). Barplots are shown for each analyzed value of K: from 2 to 8 in **E** and from 9 to 17 in **F.** (from 2 to 15). Each vertical bar represents an isolate, and the clusters are represented by different colors. Isolates are sorted by continent and by country.

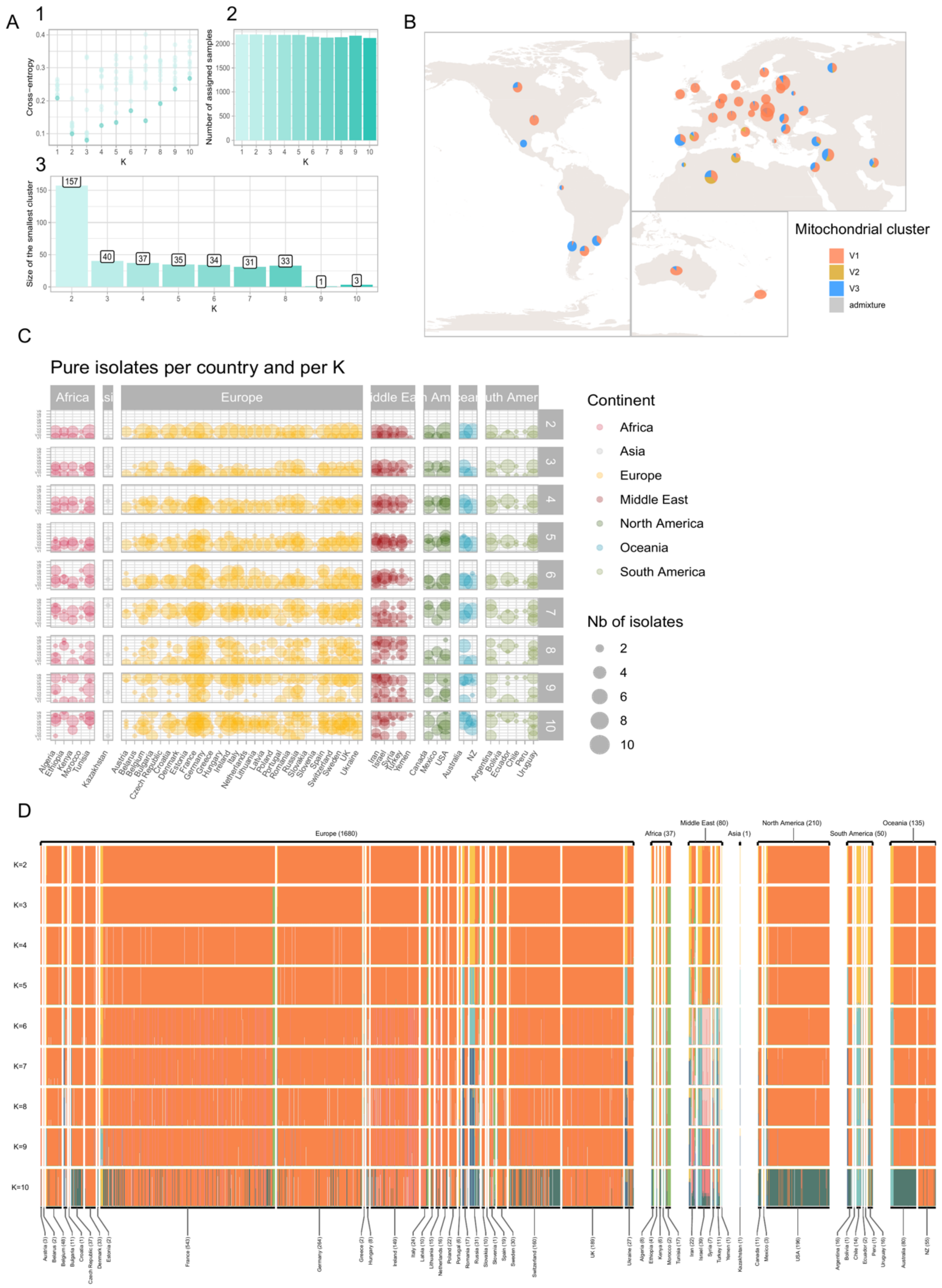


**Supplementary Figure 3**. sNMF clustering for the mitochondrial genomes, based on 17 biallelic SNPs with minor allele frequency above 0.02. **A. 1.** Cross-entropy for each number of clusters (K) as assessed by the R package *LEA*. **2.** Number of isolates uniquely assigned to one cluster per K. **3.** Number of isolates assigned to the smallest cluster. **B.** Map of the genetic clustering based on a nuclear genome SNP dataset using sNMF. Each color represents a different genetic cluster, and the sizes of the slices represent the average attribution to the cluster across the isolates from each location. The isolates defined as admixed are colored in grey. **C.** Distribution of the isolates per country per cluster for the number of clusters (K) from 2 to 10. **D.** Genetic clustering of isolates based on mitochondrial SNPs showing per-isolate barplots with cluster assignments**.** Barplots for the best replicate for each K (as determined by cross-entropy). Barplots are shown for each analyzed value of K (from 2 to 10). Each vertical bar represents an isolate, and the clusters are represented by different colors. Isolates are sorted by continent and by country.

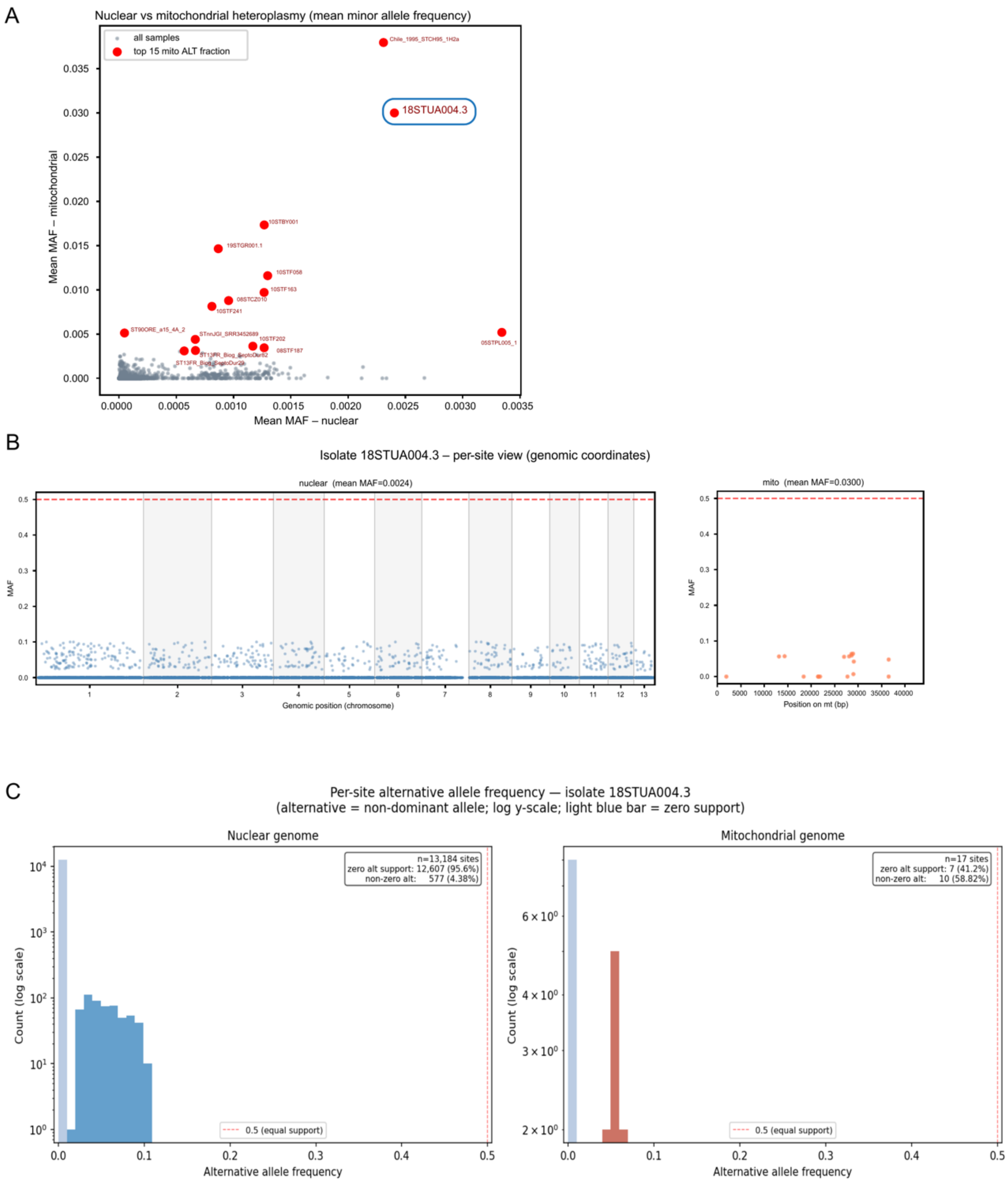


**Supplementary Figure 4.** Heteroplasmy analysis. **A.** Comparison of the mean minor allele frequency (MAF) of filtered biallelic SNPs in the nuclear and mitochondrial genomes for each isolate. The 15 isolates with the highest ratio of mean mitochondrial to mean nuclear MAF are highlighted in red and labeled. **B.** Minor allele frequencies of mitochondrial and nuclear SNPs plotted across genomic coordinates for the candidate isolate 18STUA004.3. Each point represents a polymorphic site. **C.** Distribution of mitochondrial and nuclear SNP minor allele frequencies for candidate isolate 18STUA004.3.

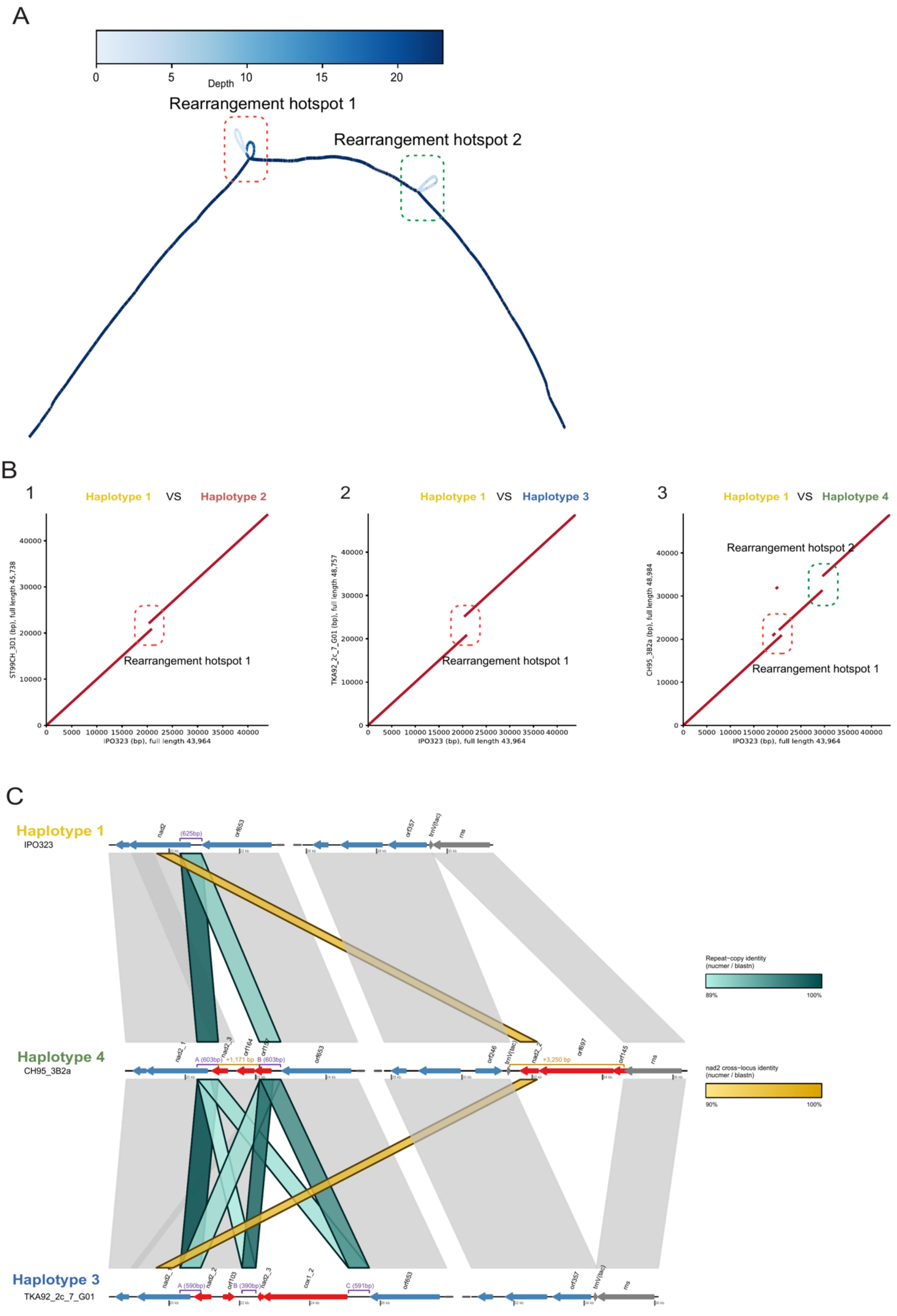
A
Depth
0
5
10
15
20
Rearrangement hotspot 1
Rearrangement hotspot 2
B
1
Haplotype 1 VS Haplotype 2
ST99CH_3D1 (bp), full length 45,738
IPO323 (bp), full length 43,964
Rearrangement hotspot 1
2
Haplotype 1 VS Haplotype 3
TKA92_2c_7_G01 (bp), full length 48,757
IPO323 (bp), full length 43,964
Rearrangement hotspot 1
3
Haplotype 1 VS Haplotype 4
CH95_3B2a (bp), full length 48,984
IPO323 (bp), full length 43,964
Rearrangement hotspot 2
Rearrangement hotspot 1
C
Haplotype 1
IPO323
Haplotype 4
CH95_3B2a
Haplotype 3
TKA92_2c_7_G01
Repeat-copy identity (nucmer / blastn)
89%
100%
nad2 cross-locus identity (nucmer / blastn)
90%
100%

**Supplementary Figure 5**. **A.** Pangenome graph of 23 PacBio-sequenced isolates. The intensity of the blue color reflects node coverage in the pangenome graph, with darker shades indicating nodes present in a larger number of genomes. The loops formed by the structural variation are framed in red (hotspot 1) and green (hotspot 2). **B.** Whole mitogenome pairwise alignments between Haplotype 1 (isolate IPO323) and **(1)** Haplotype 2 (isolate ST99CH_3D1), **(2)** Haplotype 3 (TKA92_2c_7_G01), and **(3)** Haplotype 4 (CH95_3B2a). The first and second rearrangement hotspots are marked in red and green, respectively. **C.** First and second hotspot synteny across three haplotypes on their own native coordinates, joined by a dashed line where the intervening core sequence is compressed out. Haplotype 1 (IPO323) carries neither insertion; Haplotype 4 (CH95_3B2a) carries both: a short insertion at hotspot 1 and an insertion at hotspot 2; Haplotype 3 (TKA92_2c_7_G01) carries only the large hotspot-1 insertion. Genes are shown as arrows colored by class: blue, protein-coding genes surrounding the insertions; grey, tRNAs/rRNAs; red, genes carried by an insertion. Purple brackets mark each flanking direct-repeat copy (lettered A/B/C) with its length; gold brackets give the insertion's total size. Cyan ribbons show pairwise nucleotide identity between repeat copies; gold ribbons show nucleotide identity linking nad2-derived fragments across the two hotspot loci.

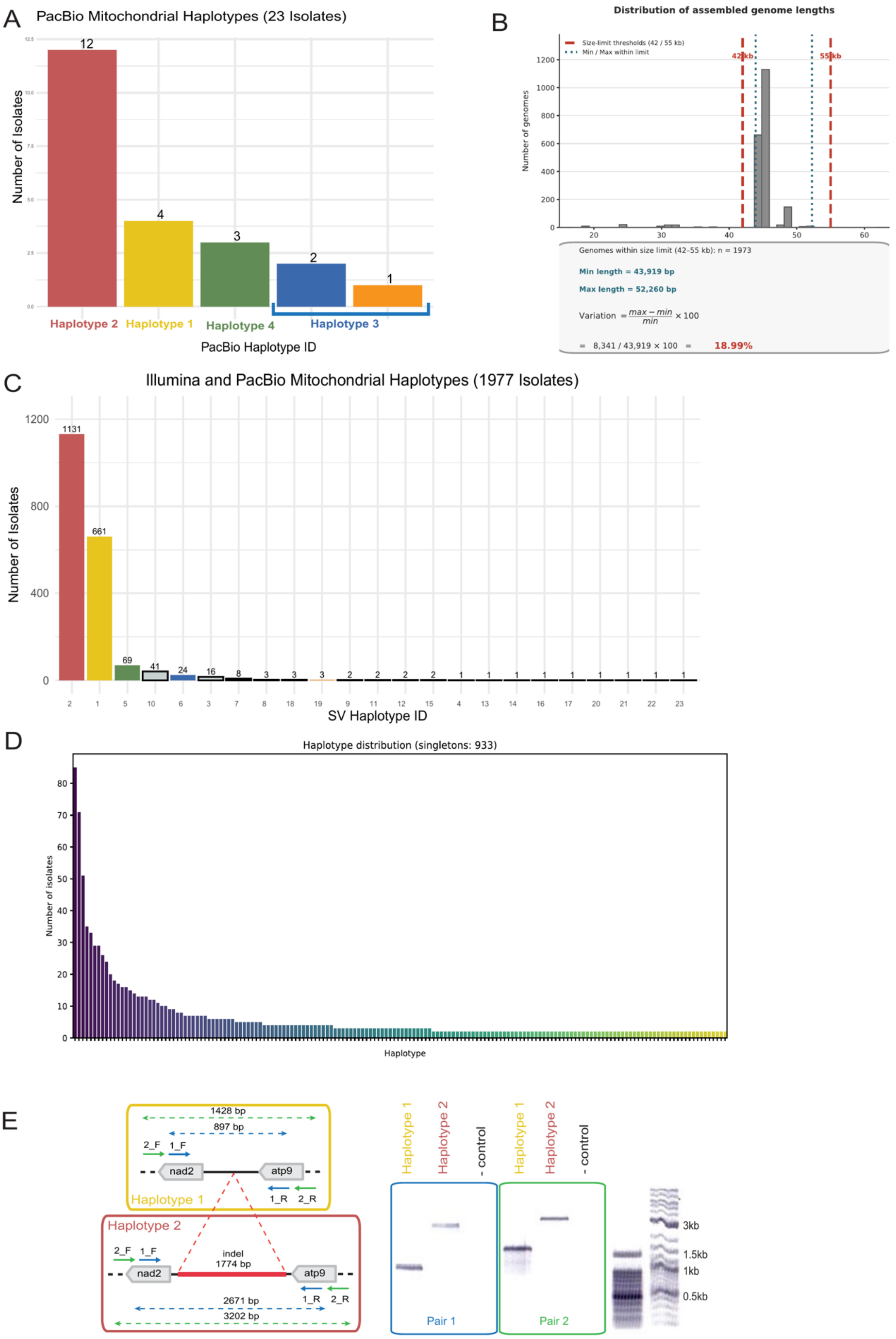

A
PacBio Mitochondrial Haplotypes (23 Isolates)
Number of Isolates
12
4
3
2
1
Haplotype 2
Haplotype 1
Haplotype 4
Haplotype 3
PacBio Haplotype ID
B
Distribution of assembled genome lengths
Size-limit thresholds (42 / 55 kb)
Min / Max within limit
42 kb
55 kb
Number of genomes
Genomes within size limit (42-55 kb): n = 1973
Min length = 43,919 bp
Max length = 52,260 bp
Variation = max − min / min × 100
= 8,341 / 43,919 × 100 = 18.99%
C
Illumina and PacBio Mitochondrial Haplotypes (1977 Isolates)
Number of Isolates
1131
661
69
41
24
16
8
3
3
3
2
2
2
2
1
1
1
1
1
1
1
1
1
2
1
5
10
6
3
7
8
18
19
9
11
12
15
4
13
14
16
17
20
21
22
23
SV Haplotype ID
D
Haplotype distribution (singletons: 933)
Number of isolates
Haplotype
E
1428 bp
897 bp
2_F
1_F
nad2
atp9
1_R
2_R
Haplotype 1
Haplotype 2
indel
1774 bp
2671 bp
3202 bp
Haplotype 1
Haplotype 2
- control
Pair 1
Pair 2
3kb
1.5kb
1kb
0.5kb

F

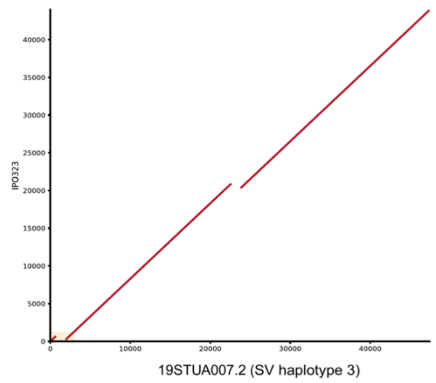


G

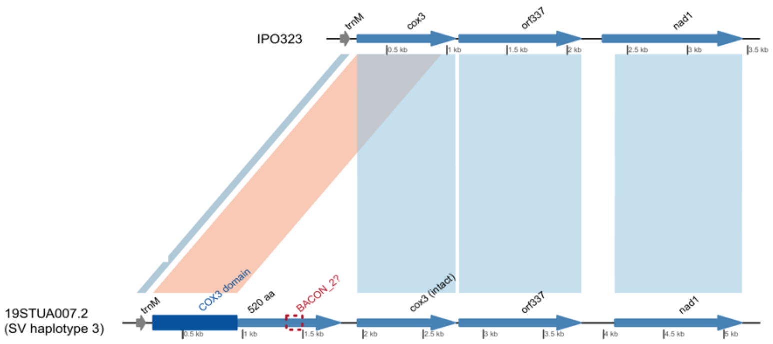


**Supplementary Figure 6**. Mitochondrial haplotypes defined based on the structural variants. **A.** Structural haplotypes were defined for the 23 PacBio-sequenced isolates. Coloring and the numbering of the haplotypes correspond to Figure 4. The fourth haplotype can be split into two if we consider the short indel of 38 bp located at position 23101 of the reference mitogenome, between the nad2 and atp9 genes. **B.** Distribution of assembled genome lengths. The red dashed lines at 42 and 55 kb mark the size limits used to keep complete assemblies. Of the genomes within these limits (n = 1,973), the shortest is 43,919 bp and the longest is 52,260 bp (dotted lines); the lengths vary by 18.99%. **C.** All structural haplotypes (based only on SVs) are defined for the 1977 PacBio- and Illumina-sequenced isolates. Coloring of the haplotypes corresponds to Figure 4. **D.** Haplotypes (based on a total of 1941 genomic variants, comprising 1178 SNPs, 745 short indels, and 18 SVs) defined for the 1977 PacBio- and Illumina-sequenced isolates by any variants. Most of the haplotypes defined by these variants are represented by only one isolate (933 out of 1099, 85%, not represented on the barplot); meanwhile, three haplotypes have more than 50 identical mitochondrial assemblies each, representing in total 10% (204 isolates) of the analyzed dataset. **E.** PCR validation of the insertion between *nad2* and *atp9*. (**Left**) Diagram of the *nad2–atp9* intergenic region in Haplotype 1 (IPO323, reference, without the insertion) and Haplotype 2 (ST95CH_3B2a, with the ~1.7 kb insertion, red). Arrows show where the two independent primer pairs bind: pair 1 (1_F/1_R, blue) and pair 2 (2_F/2_R, green). Dashed lines give the expected amplicon sizes. Pair 1 should yield 897 bp for Haplotype 1 and 2,671 bp for Haplotype 2; pair 2 should yield 1,428 bp and 3,202 bp. (**Right**) Agarose gel (0.8%) of the PCR products from both primer pairs, with molecular size markers on the right. For both pairs, Haplotype 2 gives a longer product than Haplotype 1, and each band matches its expected size, confirming that the insertion is present in ST95CH_3B2a and absent in IPO323. The negative (no-template) controls show no amplification. Detailed depiction of the structural variant from Figure 4 E in **F, G**. **F.** The indel is present in 16 isolates, always alongside the short 1767 bp insertion at the first rearrangement hotspot (PacBio-based haplotype 2), with all 16 isolates forming a distinct haplotype (haplotype 3). The SV is characterized by a 1275 bp insertion at position 231 of the IPO323 genome; the insertion is depicted in a yellow frame. **G.** This indel is identical across all 16 isolates and contains a 520-aa-long ORF with a cox3 domain and a BACON_2 domain. The insertion occurs within the coding sequence of the *cox3* gene, but it does not result in a loss of function. The flanking repeats suggest a potential duplication of the neighboring region, including the start of the *cox3* gene.

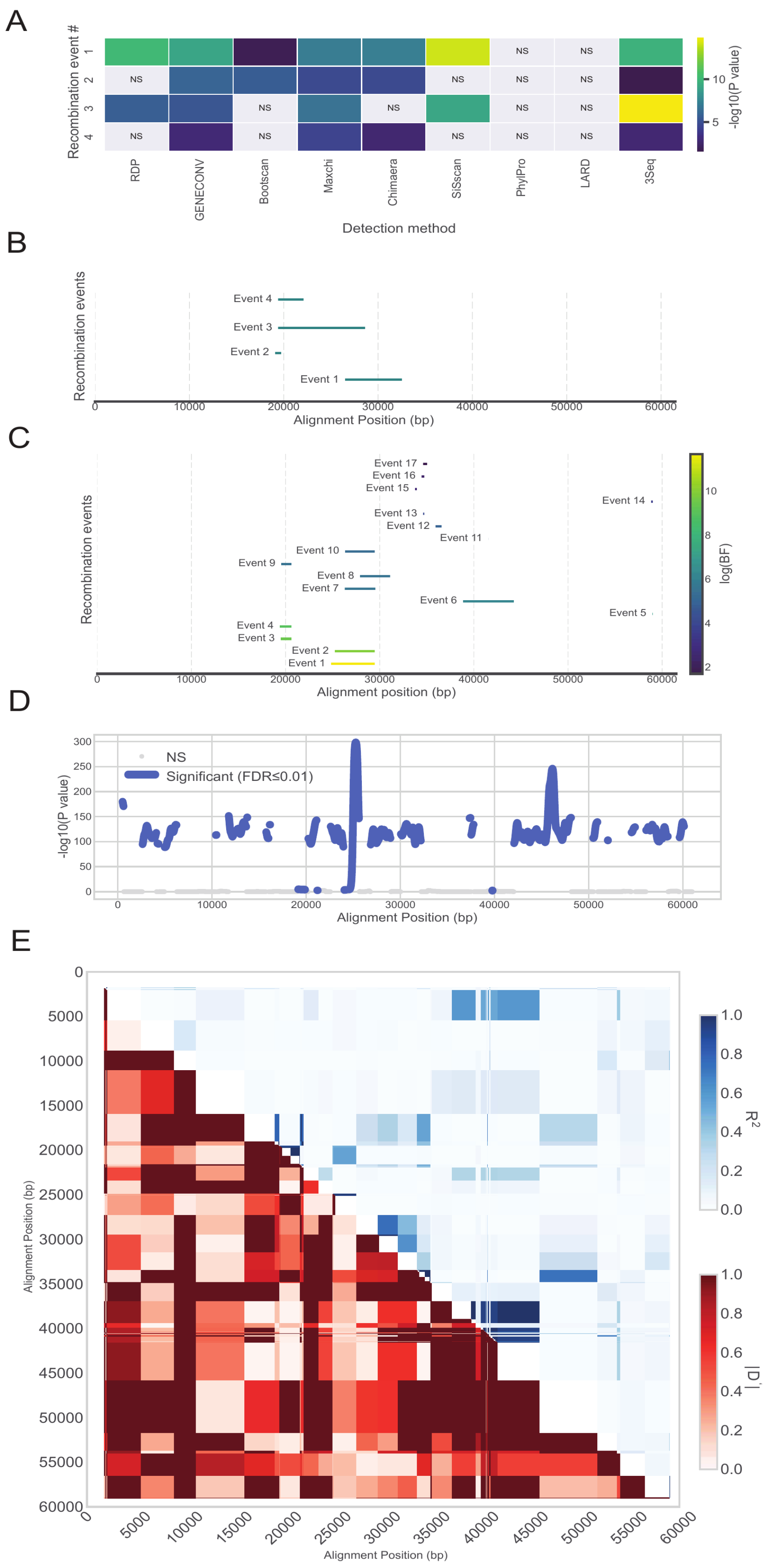
A
Recombination event #
Detection method
RDP
GENECONV
Bootscan
Maxchi
Chimaera
SiSscan
PhylPro
LARD
3Seq
NS
-log10(P value)
B
Recombination events
Event 1
Event 2
Event 3
Event 4
Alignment Position (bp)
C
Recombination events
Event 1
Event 2
Event 3
Event 4
Event 5
Event 6
Event 7
Event 8
Event 9
Event 10
Event 11
Event 12
Event 13
Event 14
Event 15
Event 16
Event 17
log(BF)
Alignment position (bp)
D
-log10(P value)
NS
Significant (FDR≤0.01)
Alignment Position (bp)
E
Alignment Position (bp)
R2
|D'|
Alignment Position (bp)

F

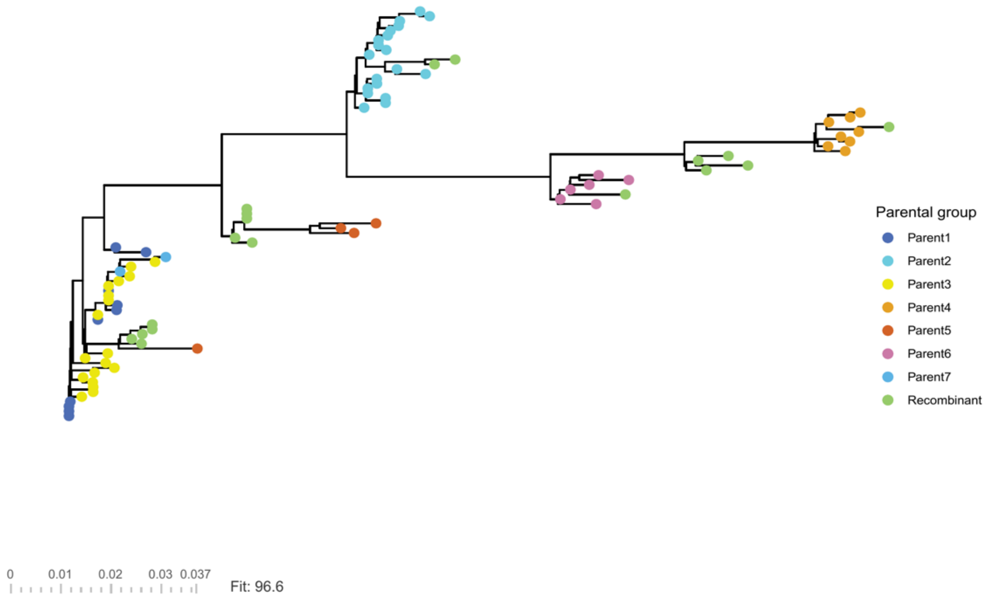


G

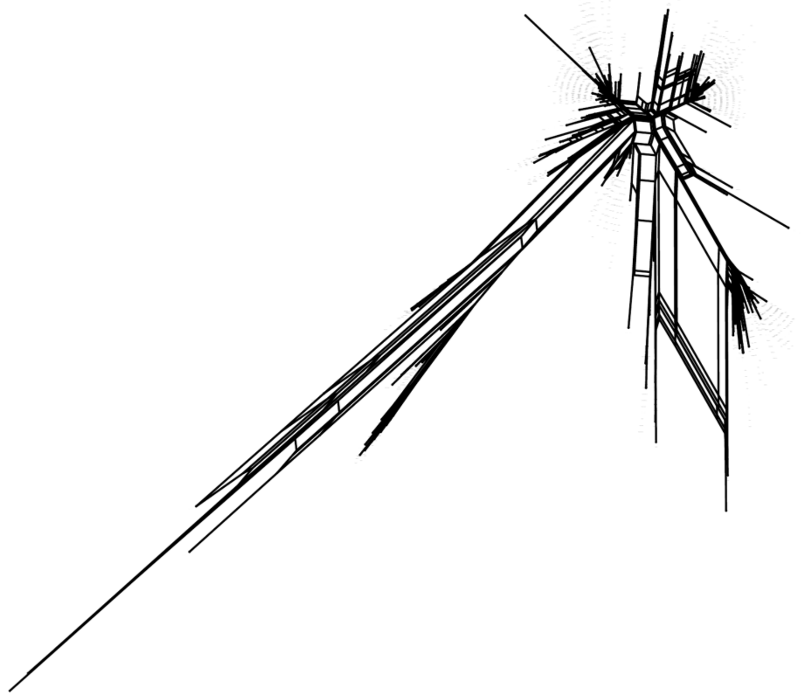

**Supplementary Figure 7**. Recombination detection. Recombination detection tools report multiple plausible recombination events in the *Z. tritici* population. **A** and **B.** Four recombination events were reported by the RDP4 methods in the full population alignment. **A** shows the -log10 P value of the identified events. GENECONV, MaxChi, and 3Seq report statistically significant recombination in all cases. Chimaera does not report the third event as recombination. RDP and Bootscan do not report two events as recombination, whereas PhylPro and LARD do not report any of the cases as recombination events. **B** shows the genomic locations of the four events. The positions of the recombination events are shown relative to the alignment and span between 20 kb and 34 kb. **C.** Recombination events identified with fastGEAR. Positions are shown relative to the alignment. The color represents the natural logarithm of the Bayes factor reported by the tool. The positions of the recombination events span between 19 kb and 45 kb. **D.** Regions with significant recombination events within a 100 bp window were identified with the PhiPack Profile tool. The FDR correction was applied to the output of the tool, and events with FDR $\leq 0.01$ are shown in blue. The two most prominent peaks are located at 25 kb and 46 kb. **E.** A heatmap of $R^2$ (in blue) and |D'| (in red) alignment-wise pairwise linkage disequilibrium (LD) measurements for SNPs. Linked alleles have values closer to 1. **F.** A phylogenetic tree constructed from the selected SNPs, as in Figure 3 D2, and painted with assigned colors. Recombinants unassigned to any of the seven parental clusters are shown in green. **G.** NeighborNet split network built in SplitsTree (fit = 96.6%). Box-like reticulations between clusters indicate conflicting phylogenetic signal consistent with recombination. Scale bar: substitutions per site.

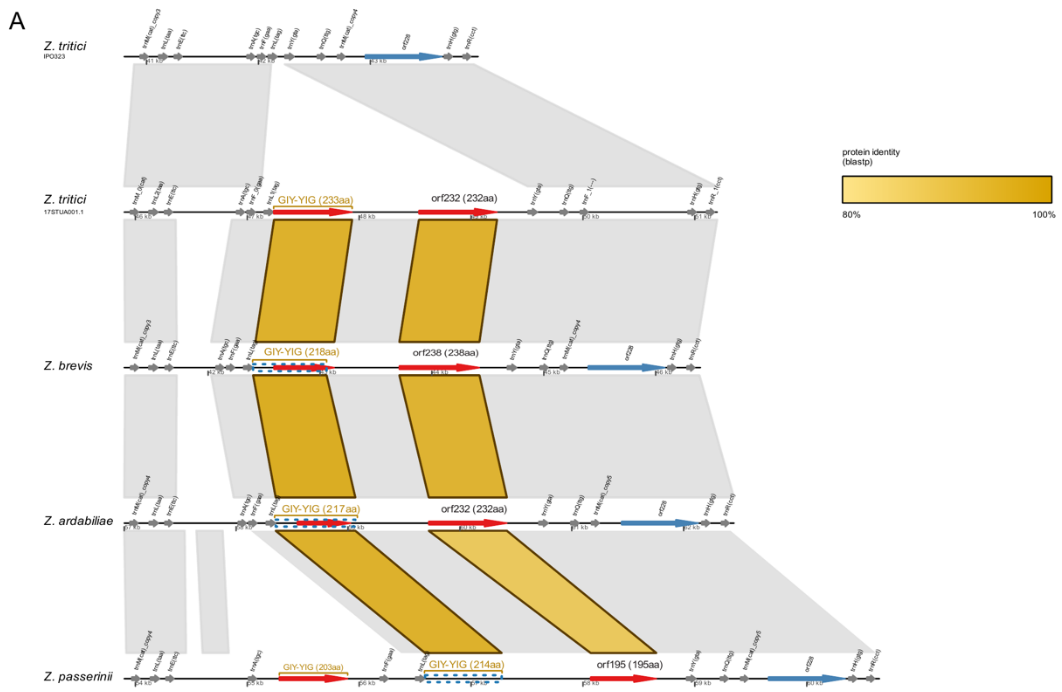


**Supplementary Figure 8**. Synteny of the mitochondrial GIY-YIG hotspot in *Zymoseptoria tritici* and three related species. Tracks from top to bottom: reference mitogenome of *Z. tritici* IPO323, *Z. tritici* 17STUA001.1 (containing the insertion with the GIY-YIG homing endonuclease), *Z. brevis, Z. ardabiliae,* and *Z. passerinii*. The gene coloring scheme is the same as in Figure 6, with the red arrows depicting the ORFs of the insertions, predicted by MFannot. These include the GIY-YIG homing-endonuclease ORFs, labeled in yellow, and a second ORF family (orf232/orf238/orf195), encoding the type I restriction enzyme R protein N-terminal domain (*HSDR_N,* Pfam PF13588). Blue dashed boxes mark the GIY-YIG sequences used in the phylogenetic analysis, recovered by blastp with a GIY-YIG from Z. tritici as the query. Grey blocks show collinear regions identified by blastn. Gold ribbons show blastp protein identity between the inserted ORFs.
